# omnideconv: a unifying framework for using and benchmarking single-cell-informed deconvolution of bulk RNA-seq data

**DOI:** 10.1101/2024.06.10.598226

**Authors:** Alexander Dietrich, Lorenzo Merotto, Konstantin Pelz, Bernhard Eder, Constantin Zackl, Katharina Reinisch, Frank Edenhofer, Federico Marini, Gregor Sturm, Markus List, Francesca Finotello

**Affiliations:** Data Science in Systems Biology, TUM School of Life Sciences, Technical University of Munich, 85354 Freising, Germany; Department of Molecular Biology, Digital Science Center (DiSC), University of Innsbruck, 6020 Innsbruck, Austria; Institute for Informatics, Ludwig-Maximilians-Universität München, 80333 München, Germany; Department of Molecular Biology, Center for Molecular Biosciences Innsbruck (CMBI), University of Innsbruck, 6020 Innsbruck, Austria; Institute of Medical Biostatistics, Epidemiology and Informatics (IMBEI), University Medical Center of the Johannes Gutenberg University Mainz, 55131 Mainz, Germany; Research Center for Immunotherapy (FZI), 55131 Mainz, Germany; Biocenter, Institute of Bioinformatics, Medical University of Innsbruck, 6020 Innsbruck, Austria; Boehringer Ingelheim International Pharma GmbH & Co KG, 88397 Biberach, Germany; Munich Data Science Institute (MDSI), Technical University of Munich, 85748 Garching, Germany

**Author notes:** **Corresponding authors:** Markus List, Francesca Finotello. Equal contribution (Alexander Dietrich, Lorenzo Merotto). Equal contribution (Markus List, Francesca Finotello).

**Keywords:** Cell-type deconvolution, transcriptomics, single-cell RNA-seq, bulk RNA-seq, method benchmark, unified method access, validation datasets

## Abstract

**Background:** In silico cell-type deconvolution from bulk transcriptomics data is a powerful technique to gain insights into the cellular composition of complex tissues. While first-generation methods used precomputed expression signatures covering limited cell types and tissues, second-generation tools use single-cell RNA sequencing data to build custom signatures for deconvoluting arbitrary cell types, tissues, and organisms. This flexibility poses significant challenges in assessing their deconvolution performance.

**Results:** Here, we comprehensively benchmark second-generation tools, disentangling different sources of variation and bias using a diverse panel of real and simulated data. Our results reveal substantial differences in accuracy, scalability, and robustness across methods, depending on factors such as cell-type similarity, reference composition, and dataset origin.

**Conclusions.:** Our study highlights the strengths, limitations, and complementarity of state-of-the-art tools, shedding light on how different data characteristics and confounders impact deconvolution performance. We provide the scientific community with an ecosystem of tools and resources, *omnideconv*, simplifying the application, benchmarking, and optimization of deconvolution methods.

## Background

Tissues and organs are composed of various cell types, collectively determining their structure and function. Therefore, characterizing the cellular composition of tissues is essential for studying cell development, homeostasis, disease, and response to therapeutic interventions. In recent years, several in silico deconvolution methods have been developed to estimate the cellular composition of tissue samples profiled with bulk RNA sequencing (RNA-seq). Deconvolution algorithms consider gene expression profiles of a heterogeneous sample as the weighted sum of the gene expression profiles of the admixed cells, and estimate the unknown cell fractions leveraging cell-type-specific transcriptomic signatures [1]. While single-cell RNA-seq (scRNA-seq) enables studying the transcriptomes underlying cellular identities at unprecedented resolution and granularity [2], it is not suited to accurately quantify the cellular composition of tissues. This is mainly due to differences in single-cell dissociation efficiency, which can bias cell-type proportions [3–6]. Moreover, single-cell protocols entail considerable costs and technical challenges, making their application unattractive for profiling extensive sample collections. Thus, bulk transcriptome profiling remains popular, motivating further research into in silico cell-type deconvolution.

The *first generation* of deconvolution methods mainly relied on bulk transcriptomic data from purified/enriched cell types to derive cell-type-specific transcriptional “fingerprints”, which were then directly embedded into the tools and usable as precomputed signature matrices. First-generation methods only covered a limited set of cell types for which purified transcriptomic data were available (i.e., mainly from human immune cells, with a few exceptions) [7], but had the advantage of being extensively validated [8–12]. The need to extend deconvolution analyses to underrepresented cell types and tissues has prompted some first-generation tools to support user-defined signature matrices [13–17] or to enhance precomputed signatures by incorporating profiles derived from scRNA-seq data [14]. Building upon these pioneering efforts and further leveraging the rapid evolution of the single-cell transcriptomics field, a *second generation* of deconvolution tools has emerged [18]. These tools directly learn cell-type-specific signatures from annotated (i.e., cell-type-labeled) scRNA-seq data, allowing, in principle, the deconvolution of any cell type, tissue, and organism, for which single-cell data is available. As second-generation methods flexibly derive deconvolution signatures “on the fly”, depending on the user-specified data, characterizing their accuracy and robustness in different contexts requires systematic and comprehensive benchmarking that differs from previous studies focused on first-generation methods. While some second-generation algorithms have been previously tested [5,19–25], major challenges in deconvolution benchmarking remain unaddressed [6,25,26]. These include assessing the methods’ ability to quantify rare or closely-related cell types, and determining the impact of biological and technical biases on deconvolution performance.

In this study, we comprehensively benchmark second-generation deconvolution tools, leveraging a balanced and rationally-designed set of simulated and experimental ground-truth data, while ensuring reproducibility and reusability. To disentangle and systematically assess the impact of various biological and technical confounders on methods performance, we used our previously developed simulator SimBu [27], which allows for the efficient generation of synthetic bulk RNA-seq datasets, i.e., ‘pseudo-bulks’ generated by the controlled aggregation of single-cell expression profiles. SimBu enables the modeling of cell-type-specific mRNA content, a key source of bias that deconvolution methods have to account for [1]. We complemented our set of pseudo-bulk data with real RNA-seq samples from different tissues and organisms with matching ground-truth cell fractions. Overall, we assembled a compendium of more than 5,500 real and simulated RNA-seq samples and matched ground-truth cell fractions (**Table S1**) to systematically test method performance in different scenarios.

Benchmarking studies typically represent a snapshot of available methods in a fast-evolving field, making it challenging to consider additional tools and datasets in follow-up investigations [28]. Moreover, users might need to further evaluate and possibly optimize their deconvolution strategies on their specific use case, which might target cell types, tissues, or even organisms not covered by previous assessments. To overcome these limitations, we assembled a freely available ecosystem of tools and resources called *omnideconv* (omnideconv.org), which enables the simplified use and assessment of cell-type deconvolution methods (**Fig. 1A**). Besides SimBu and our collection of validation datasets (*deconvData*), our ecosystem includes: 1) *omnideconv,* a novel R package that offers uniform access to several second-generation deconvolution methods, 2) *deconvBench*, a Nextflow [29] pipeline to reproduce and extend the presented benchmarking study, and 3) *deconvExplorer,* a web app to interactively investigate deconvolution signatures and results. We envision that the omnideconv ecosystem will aid researchers in deconvolving RNA-seq samples more easily and offer guidance for method choice in different scenarios. In addition, the flexibility of our applications allows for easy extension, which is a necessary feature to include upcoming deconvolution methods, as well as to benchmark and optimize them for specific applications.

**Fig. 1:**
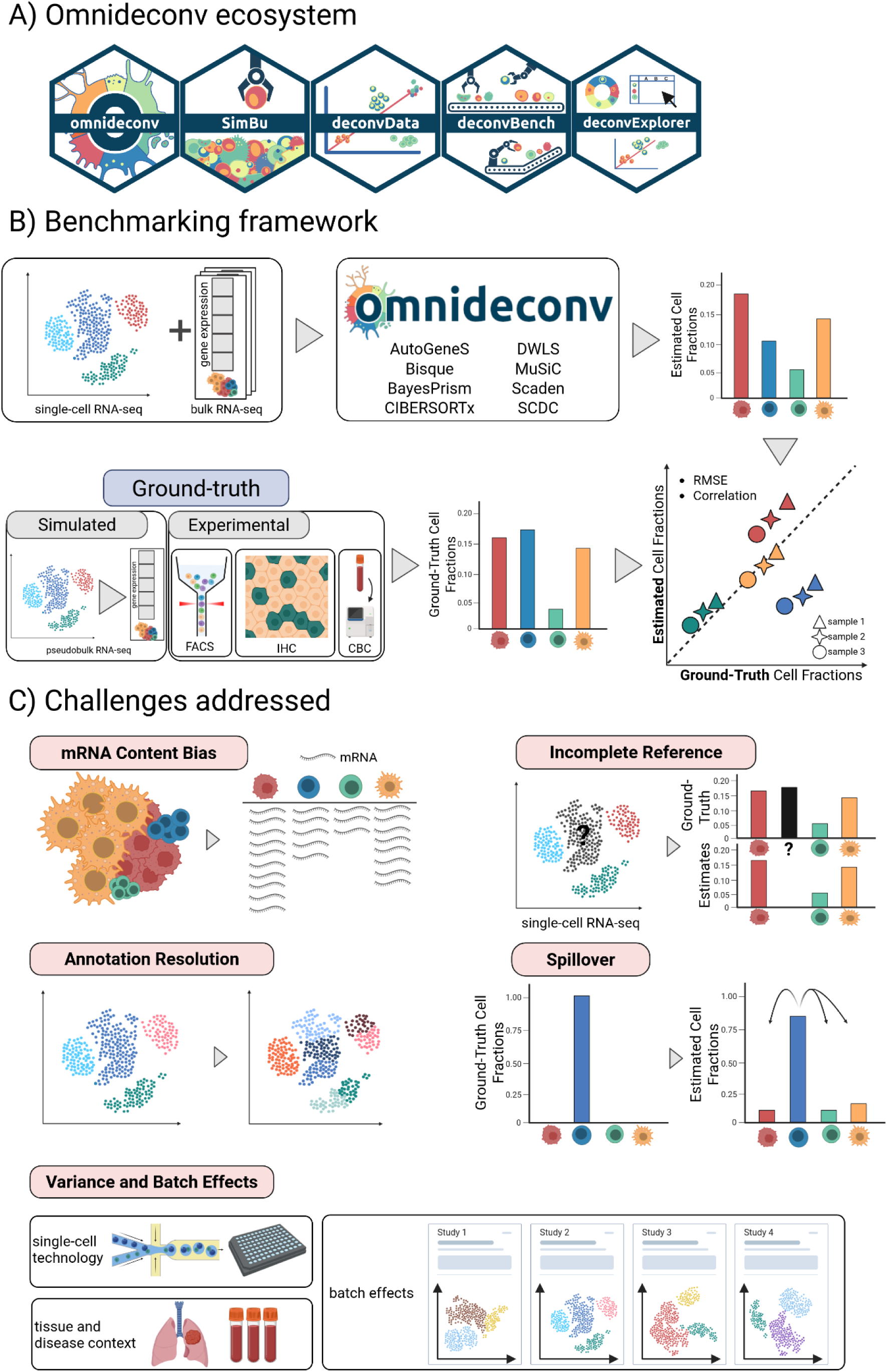
omnideconv ecosystem and benchmark **(A)** The omnideconv benchmarking ecosystem offers five tools (from left to right): the R package *omnideconv* providing a unified interface to deconvolution methods, the pseudo-bulk simulation method SimBu, the *deconvData* data repository, the *deconvBench* benchmarking pipeline in Nextflow, and the web-app *deconvExplorer*. **(B)** Outline of the benchmark experiment: scRNA-seq and bulk RNA-seq data are used as input for several methods, and a unified output of estimated cell-type fractions for each bulk sample is calculated. We compare the estimated fractions to ground-truth fractions (known from pseudo-bulks or FACS/IHC/CBC assays) and compute performance measures per method and cell type. **(C)** Several challenges in cell-type deconvolution are addressed in this benchmark: (1) cell types show total mRNA bias; (2) a fraction of the cells may be of unknown type, since the scRNA-seq reference does not necessarily contain all cell types present in the bulk mixture; (3) the level of annotation granularity; (4) some cell types are more similar on a transcriptomic level, leading to “spillover” towards similar cell types; (5) scRNA-seq datasets vary by technology, tissue and batch effects. This figure was created with Biorender.com.

## Results

### The omnideconv ecosystem enables the simplified application and benchmarking of second-generation deconvolution methods

Second-generation deconvolution methods differ in programming languages, workflows, input data types, and processing approaches. This complicates the simultaneous execution of multiple methods and the comparison of their estimates. To simplify the usage of second-generation deconvolution methods, we have developed *omnideconv* (https://github.com/omnideconv/omnideconv/), an R package providing a unified interface to multiple R- and Python-based methods. *omnideconv* currently supports twelve methods: AutoGeneS [30], BayesPrism [31], Bseq-SC [32], Bisque [33], CDseq [34], CIBERSORTx [35], CPM [36], DWLS [37], MOMF [38], MuSiC [39], SCDC [40], and Scaden [41]. It unifies the methods’ workflows, input and output data, and semantics, allowing deconvolution analysis using one or two simple commands.

For this study, we selected eight methods embedded within the omnideconv R package (AutoGeneS, BayesPrism, Bisque, CIBERSORTx, DWLS, MuSiC, SCDC, and Scaden) that: 1) leverage annotated scRNA-seq data to directly perform cell-type deconvolution (rather than reference-free deconvolution followed by a posteriori annotation of derived cell phenotypes); 2) do not strictly require context-specific marker genes; and 3) provide deconvolution results in the form of relative cell fractions. Of note, we implemented and evaluated an optimized version of DWLS in *omnideconv,* with considerably more effective usage of computational resources (“DWLS optimized”, details in Methods). In contrast to earlier studies assessing normalization and parameter optimization [20,28], we consulted the methods’ developers for guidance on optimal parameter settings and data type usage. In the absence of feedback, we ran the corresponding method with default parameters and considered counts and transcripts per million (TPM) as single-cell and bulk RNA-seq input data, respectively (details in Methods); of note, CIBERSORTx was run through the official docker image; we omitted batch correction due to suboptimal results obtained with S-mode.

Our benchmarking study (**Fig. 1B**) comprises human and mouse bulk RNA-seq datasets for which ground-truth estimates of cell fractions were available from fluorescence-activated cell sorting (FACS), immunohistochemistry (IHC), or complete blood counts (CBCs). We did not extend our set of validation datasets to those based on single-cell-derived “ground truth” cell fractions due to the well-known limitations in cell-capture efficiency of scRNA-seq technologies [3–6]. Similarly, we excluded RNA-seq datasets derived from RNA mixtures rather than cell mixtures [42], as they do not capture cell-type-specific mRNA biases [43]. We made our selection of validation datasets readily available to the scientific community through *deconvData* (https://figshare.com/projects/deconvData/197794). In addition to these real datasets, pseudo-bulk datasets were generated using SimBu [27] to systematically test methods’ performance in various scenarios and evaluate their robustness to different confounders. In particular, we assessed the impact of distinct factors that can bias cell-type estimates: 1) cell-type-specific mRNA bias, since cells with higher overall mRNA abundance can be overestimated, and vice versa; 2) unknown cellular content, i.e., cell types present in the bulk RNA-seq data to be deconvolved but not in the scRNA-seq reference used for method training; 3) the impact of the resolution of cell type annotations (i.e., from coarse to fine-grained); 4) transcriptional similarity between closely-related cell types; 5) variance and batch effects affecting reference single-cell datasets derived from different technologies, studies, and tissue/disease context (**Fig. 1C**). All the analyses performed in this benchmarking study were implemented in a well-documented, reusable, and extensible Nextflow [29] pipeline called *deconvBench* (https://github.com/omnideconv/deconvBench).

### Deconvolution performance on real and pseudo-bulk RNA-seq datasets

We evaluated methods’ performance on immune-cell-type quantification, a key task in deconvolution analysis [8,18]. We selected four blood-cell-derived bulk RNA-seq datasets (*Finotello [43]*: n=9, *Hoek [44]*: n=8, *Morandini [45]*: n=114, and *Altman* [46]: n=322; **Table S1**) for which ground-truth cell fractions were available from FACS or CBC. For method training, we selected the *Hao* CITE-seq single-cell dataset from human peripheral blood mononuclear cells [47] (PBMC) (Methods, **Fig. S1**, and **Table S2**). We fed 10% of the single-cell data to the deconvolution methods, maintaining the original cellular composition (*HaoSub* dataset, with 14,744 cells encompassing eleven cell types; details in Methods). The generated pseudo-bulk datasets mirror the cell-type composition and sequencing depth of the real bulk RNA-seq samples, while using only cell types that exist in both the reference and the bulk datasets. This collection of real and pseudo-bulk data allowed the comparison of predicted and ground truth cell-type fractions in complex and easy scenarios, respectively, and the evaluation of deconvolution performance in terms of Pearson correlation (R) and root-mean-square error (RMSE) (Methods).

On human pseudo-bulk samples, all methods exhibited high correlation and low RMSE (**Fig. S2B and S3**). Using the same single-cell dataset for simulation and signature building can be considered the simplest scenario for deconvolution, especially for methods that create pseudo-bulks to train their internal model [41]. Despite the overall good performance, Bisque showed systematic estimation bias for T cells and CD8^+^ T cells in the *HoekSim* and *FinotelloSim* datasets, respectively. At the same time, SCDC underestimated CD4^+^ T cells in the *FinotelloSim* and *MorandiniSim* datasets.

The real bulk RNA-seq datasets represented a more challenging scenario, resulting in greater variability in method performance. Scaden could consistently reach high performances in terms of *global* correlation (0.79-0.95) and RMSE (0.06-0.15) (i.e., computed considering all cell types together) in the *Hoek*, *Finotello,* and *Morandini* datasets (**Fig. 2A and S3**). Bisque and DWLS all performed similarly well on the *Hoek* and *Morandini* datasets, while their performance dropped on the *Finotello* dataset, which encompasses more fine-grained cell types, only reaching global correlation values of 0.71 and 0.42, respectively. MuSiC, SCDC, and BayesPrism had the highest RMSE values due to their systematic *estimation biases* (i.e., under- or over-estimation) of T-cell subpopulations, monocytes, and NK cells. This issue also affected other methods, such as CIBERSORTx, though to a lesser extent. AutoGeneS exhibited a dataset-specific performance similar to Bisque and DWLS, although with overall lower accuracy, particularly on the *Finotello* dataset (R = 0.08).

**Fig. 2:**
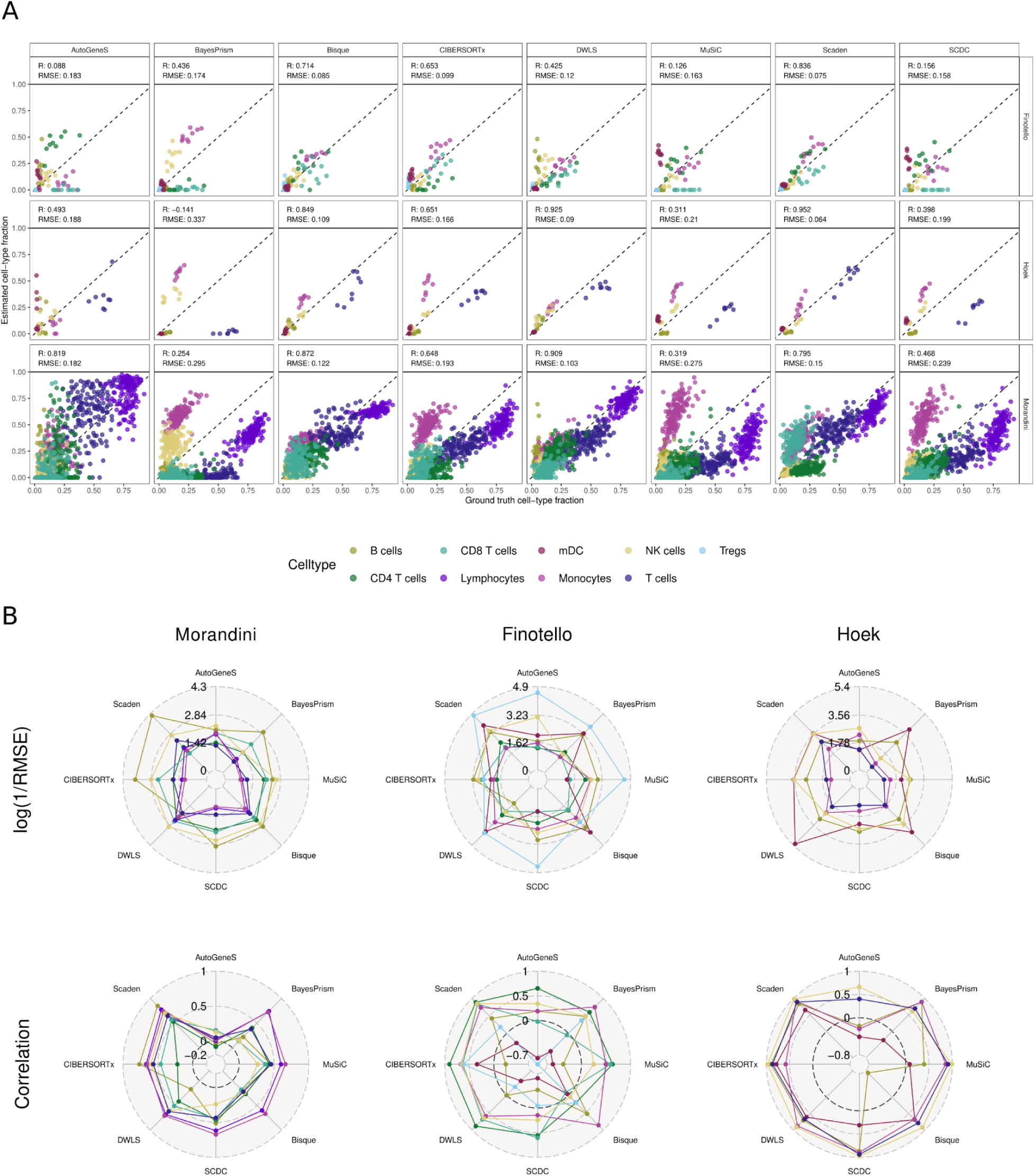
Deconvolution performances of eight second-generation methods. Predicted cell-type fractions of eight methods are compared against the ground truth fractions. The performances on three bulk (*Morandini*: n=114, *Finotello*: n=9, *Hoek*: n=8) datasets are displayed. The scRNA-seq reference dataset is *HaoSub*. (**A**) directly shows the predicted vs. estimated cell-type proportions, split by deconvolution method and dataset, and colored by cell type. A point on the dashed line represents a perfect prediction of a cell type in one sample. In (**B**), these results are aggregated into mean log(1/RMSE) and Pearson Correlation per cell type, stratified by bulk dataset (columns), and cell type (color). A datapoint in the center of the circle represents a cell type that was not detected at all by a method. A method is ideally located at the outer edge of the radar plot in both the correlation and RMSE rows, indicating high prediction quality for a cell type. Results for their pseudo-bulk representatives and three additional datasets can be found in Fig. S2-S3.

To better understand these differences, we next assessed cell-type-specific performance metrics alongside the global values (**Fig. 2B**, Methods). Cell-type-specific performance varied across methods and datasets, with some methods, such as Scaden, CIBERSORTx, and DWLS, achieving consistently good results, i.e., high correlation and low RMSE values (**Fig. 2B and S3**). Other methods showed larger estimation errors for certain cell types, particularly T-cell subpopulations, while B cells and NK cells were generally estimated more accurately across datasets. On the *Finotello* dataset, AutoGeneS, BayesPrism, MuSiC, and SCDC showed false negative predictions, i.e., failed to detect CD4⁺, CD8⁺ T cells, and B cells, highlighting the challenges in resolving low-abundance or closely related cell types. Importantly, we observed that correlation and RMSE can provide orthogonal information. For example on the *Hoek* dataset, Bisque showed a low correlation for B cells (R = -0.68) despite a low RMSE (0.05), while BayesPrism obtained high correlation but also high RMSE for monocytes due to their systematic overestimation. These examples illustrate how both metrics complement each other, especially in the presence of rare or highly-abundant cell types.

To evaluate the methods’ performance on a different organism, we considered two publicly available mouse bulk RNA-seq datasets with matched immune-cell fractions from FACS (*Chen [48]*: n=12 and *Petitprez [49]*: n=14), as well as their pseudo-bulk counterparts (*ChenSim* and *PetitprezSim*). We used a subset of the spleen Tabula Muris scRNA-seq dataset for training (*TM* with 9,083 cells, **Fig. S1**). Performance again differed substantially between simulated (for *PetitprezSim* the correlation coefficients are close to 1; only Bisque dropped to R=0.796) and real datasets (**Fig. S2, S3**), where Scaden (R=0.771, RMSE=0.164), DWLS (R=0.768, RMSE=0.166), and CIBERSORTx (R=0.752, RMSE=0.158) showed the best overall performance on the *Petitprez* dataset. Bisque also suffered from large estimation bias in both mouse-derived pseudo-bulk datasets (*ChenSim* and *PetiptrezSim*), with over-estimation of B cells and under-estimation of T-cell subtypes and monocytes.

While our study focuses on deconvolution performance in terms of agreement with ground truth estimates, the final choice of method for a specific application also depends on its computational burden. We assessed the scalability of deconvolution methods with respect to reference size and sample count (Methods, **Fig. S4**), treating the signature-creation and the deconvolution steps separately wherever possible (e.g., CIBERSORTx, DWLS, SCDC, Scaden). For a 50,000-single-cell reference and 50-sample bulk, runtimes ranged from 1 to 146 minutes, increasing to 2 to 309 minutes with a 100,000-cell reference. MuSiC was the fastest, while DWLS was the slowest in both scenarios. DWLS exhibited the steepest runtime increase with larger references (12 minutes for 1,000 cells to 309 minutes for 100,000 cells), whereas AutoGeneS showed minimal change (20 minutes for 1,000 cells, 19 minutes for 100,000 cells). For methods with separate signature matrix computation (CIBERSORTx, DWLS, Scaden), the runtime increase was primarily due to the signature creation phase. For these methods, it would be convenient to decouple these two steps, so that the signature is created only once and then reused for compatible datasets. Runtime was most sensitive to the number of samples in BayesPrism, DWLS, and AutoGeneS. BayesPrism showed the largest increase: from 10 minutes for five samples (and a 1,000-cell reference) to 240 minutes for 200 samples. The signature-creation step required the most memory, with DWLS, SCDC, Scaden, and CIBERSORTx, showing similar scaling patterns. DWLS had the highest memory demand once the reference exceeded 5,000 cells, requiring over 200 GB for a 100,000-cell reference. Scaden needed up to 100 GB. Memory usage scaled more moderately for the deconvolution step of DWLS and Scaden, remaining the lowest overall. AutoGeneS and CIBERSORTx had similar memory increases, with AutoGeneS requiring 40 GB for a 100,000-cell reference. Other methods (SCDC, MuSiC, BayesPrism, Bisque) showed a steeper memory increase, with Bisque requiring the most memory once references exceeded 10,000 cells. The number of samples had little effect on memory usage.

### Impact of reference atlas cellular resolution

Single-cell atlases can be annotated with cell labels representing coarse lineages or more fine-grained subsets and states. We considered two scRNA-seq reference datasets from human lung cancer (*Lambrechts [3]*) and breast cancer (*Wu* [50]), as well as from the Allen Brain Atlas (*Allen* [51]), to test the impact of varying cellular resolutions on deconvolution performance (Methods, **Tables S2 and S3**).

On the tumor datasets, DWLS and MuSiC showed the most robust performance across all resolutions (**Fig. 3A and S5**), as their RMSE and correlation values remained in a similar range (R > 0.75, RMSE < 0.1) for almost all cell types. DWLS obtained the best results on fine-grained deconvolution, followed by CIBERSORTx and Scaden. Bisque showed the weakest performance on fine resolution and the most marked performance increase towards coarse annotations. Conversely, BayesPrism and AutoGeneS showed decreased performance when trained with coarser cell types, especially for T and NK cells. Interestingly, AutoGeneS, Scaden, and CIBERSORTx benefited from running deconvolution at a more fine-grained resolution and subsequently aggregating deconvolution results into coarse cell types (**Fig. 3B and S6**), suggesting that this agglomerative strategy could be routinely considered for these methods.

**Fig. 3:**
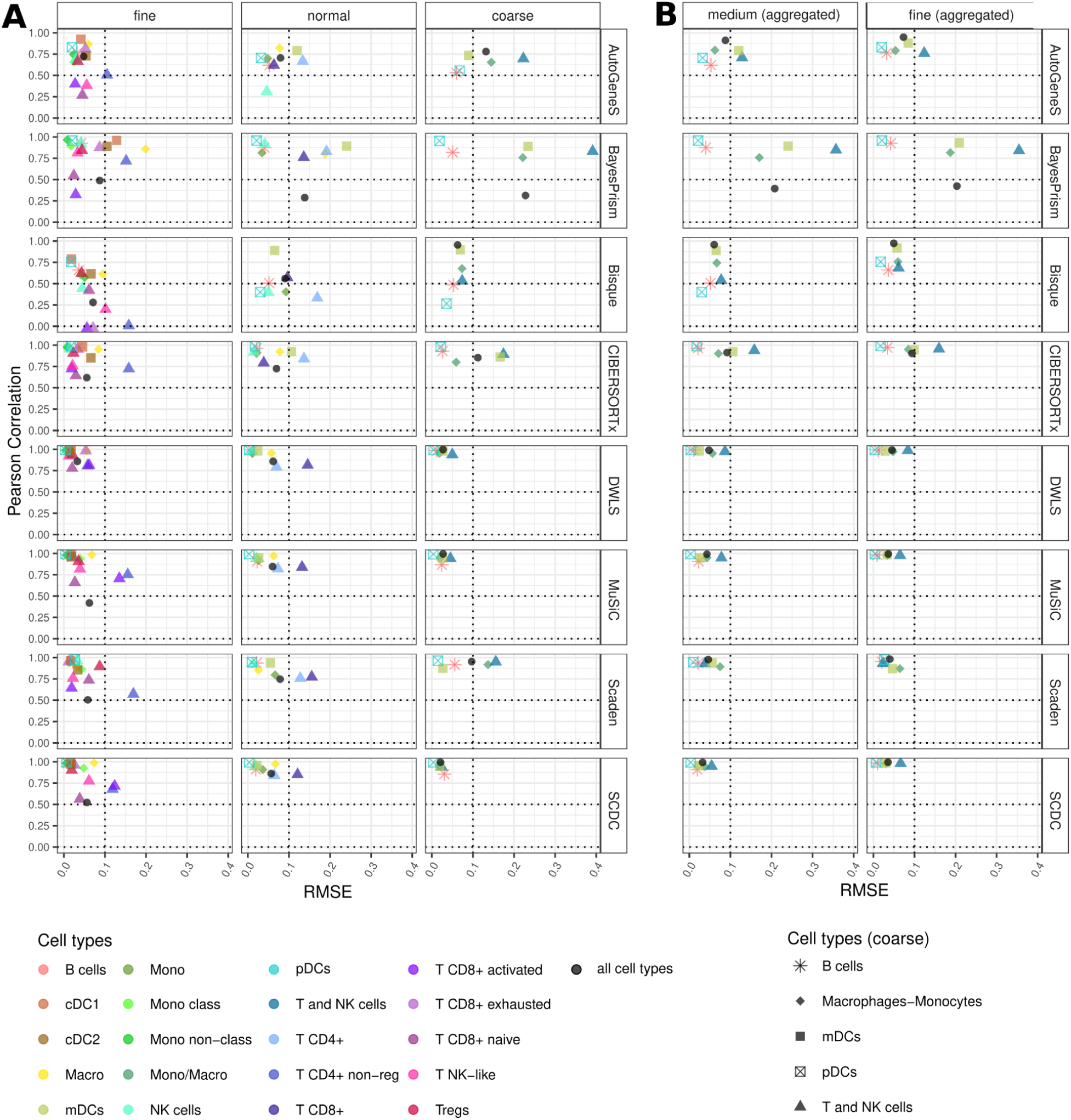
Method performances with different annotation granularities of the reference; *Lambrechts* dataset. **(A)** Pearson correlation coefficient and RMSE values computed for the cell-type-specific estimates obtained on pseudo-bulks (n=50) simulated from the *Lambrechts* dataset. **(B)** Cell-type fractions at finer resolution levels (fine, normal) were aggregated to obtain “coarse” estimates (Methods, Table S3). Cell type abbreviations: CD4^+^ T cells non-regulatory (T CD4+ non-reg), conventional dendritic cells 1 (cDC1), conventional dendritic cells 2 (cDC2), Macrophages (Macro), myeloid dendritic cells (mDCs), Monocytes (Mono), Monocytes classical (Mono class), Monocytes non-classical (Mono non-class), Monocytes/Macrophages (Mono/Macro), natural killer cells (NK cells), T cells NK-like (T NK-like), plasmacytoid dendritic cells (pDCs), T regulatory cells (Tregs).

The *Allen* brain dataset offers a unique opportunity to assess method performance on fine-grained annotations for inhibitory and excitatory neurons, which have been classified into six and eight subsets, respectively (**Table S3**). Thus, we specifically focused on these two cell types for this dataset. Overall, subsets of inhibitory neurons showed better estimates than the excitatory ones, with lower RMSE for every method. DWLS, SCDC, and MuSiC, followed by BayesPrism and CIBERSORTx, obtained the best estimates for these fine-grained cell types (i.e., high correlation and low RMSE, **Fig. S6A**). By comparing the results obtained in the direct deconvolution of the coarse cell types vs. a posteriori aggregation of the fine-grained estimates (**Fig. S6B**), we can see that certain methods again improved their deconvolution performance leveraging the “aggregation strategy”: AutogeneS, Bisque, CIBERSORTx (both cell types), and DWLS, MuSiC, and BayesPrism (inhibitory neurons only).

### Sources of systematic bias

We have seen that methods can systematically over- and under-estimate specific cell types **(Fig. 2 and S2-4**). We investigated three aspects that potentially contribute to this bias: cell-type-specific mRNA content, unknown cellular content in the (pseudo-)bulk RNA-seq data, and the transcriptional similarity of closely-related cell types. Cell types differ in their gene expression profiles but also in their size and metabolic activity [9], resulting in differences in the amount of mRNA they synthesize [52]. Methods not properly accounting for this cell-type-specific property may be prone to systematic estimation bias [1,6]. We used SimBu to create pseudo-bulk samples from eight different datasets of circulating immune cells (**Table S2**), both with and without cell-type-specific mRNA bias. As we previously showed [27], this bias can be modeled using the number of expressed genes per cell (**Fig. S1**).

By computing the difference between the RMSE on the datasets simulated with bias and without bias, we can see two main categories of methods emerging (**Fig. 4**). DWLS, Scaden, SCDC, and MuSiC improved their performance (i.e., obtained a lower RMSE) on the realistic datasets affected by cell-type-specific mRNA bias compared to their simplistic counterparts with no mRNA bias (green cells in the heatmap). Vice versa, CIBERSORTx, AutogeneS, and BayesPrism show a degraded performance when the bias is added (purple cells). Finally, Bisque is almost unaffected by the presence/absence of mRNA bias. These method-specific differences are visible for all datasets, although sometimes with more heterogeneous patterns. We further corroborated these results using the *Allen* dataset, which is suited for this investigation due to the marked differences in cell size and mRNA abundance characterizing the different cell types in the human brain [53]. The most affected cell types were those with a particularly low number of expressed genes (i.e., low mRNA content) in the single-cell data, such as platelets and erythrocytes, or vice versa, high mRNA content, such as dendritic, plasma cells, and excitatory/inhibitory neurons (**Fig. S1**). Innate lymphoid cells (ILCs) and neutrophils, which are unrepresented in the scRNA-seq reference, showed instead more variable results, likely because the mRNA-dependent error was entangled with the deconvolution-specific one.

**Fig. 4:**
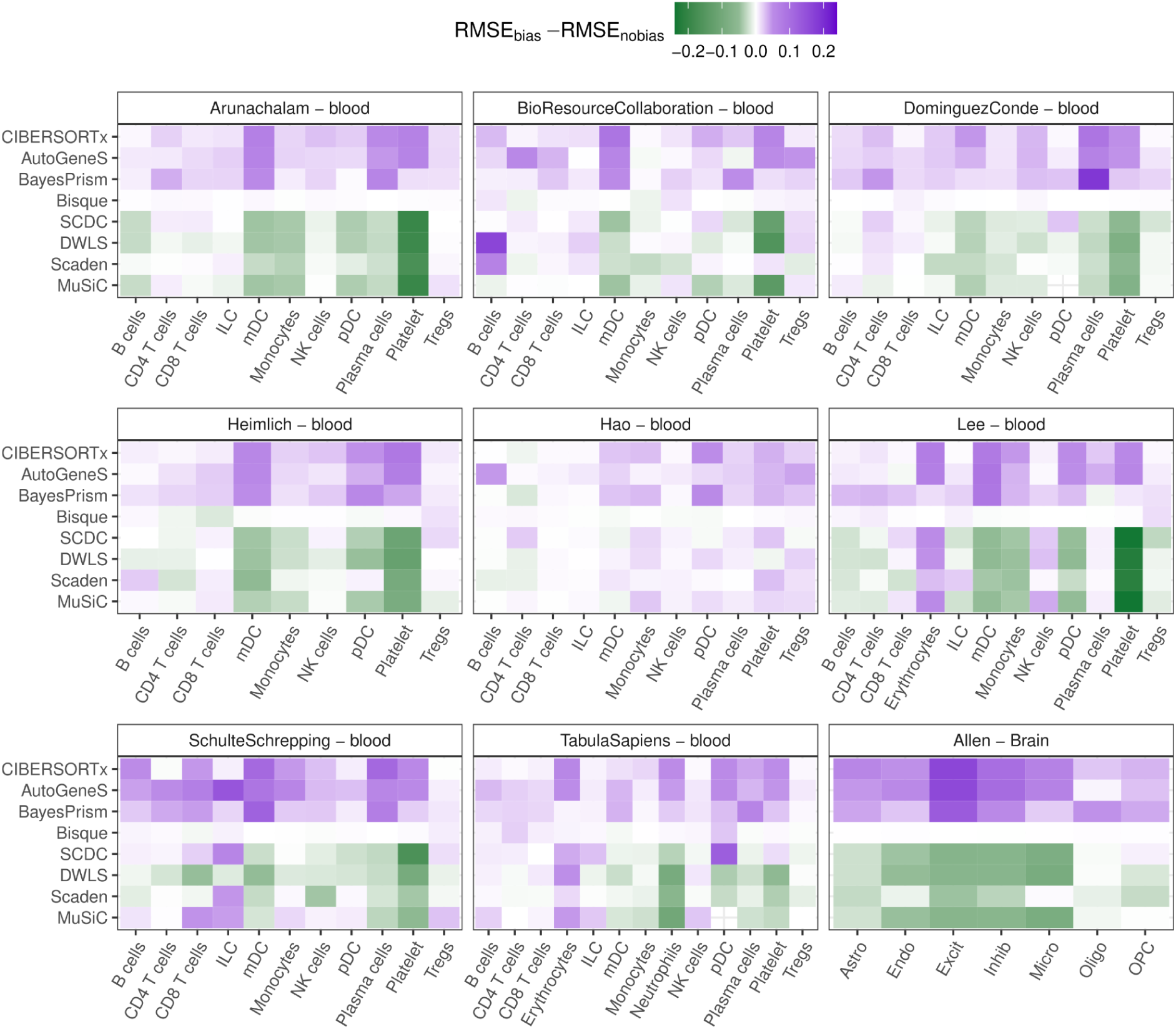
RMSE value differences between samples simulated with and without mRNA bias. Starting from eight blood and one post-mortem brain scRNA-seq dataset, we simulated pseudo bulk RNA-seq samples with and without mRNA bias. For each of those, we computed the RMSE between ground-truth and predicted cell fractions, and then the difference between the RMSE of the dataset with and without bias. A positive (purple) value indicates that the RMSE is higher in samples with bias, suggesting a worse performance with real bulk RNA-seq samples. A negative (green) case shows that the RMSE is lower when the bias is taken into account.

All the methods evaluated in this benchmarking assume that the provided single-cell dataset is an exhaustive reference of the cell types present in bulk RNA-seq to be deconvolved, and constrain the estimated cell fractions to sum up to 100% of the sample. This is rarely the case in real applications because publicly available single-cell datasets might not cover all the cell subsets present in the bulk RNA-seq data under investigation. The absence of a cell type from the training step might result in inaccurate signatures and estimation bias. To assess this, we repeatedly deconvolved the *Finotello* and *Hoek* datasets, each time removing one cell type from the *HaoSub* reference used for method training. We quantified the impact by calculating the difference in RMSE (**Fig. S7**) obtained with the “incomplete” vs. the full reference. The removal of one cell type resulted in cell-type-specific estimation bias for all methods, although to varying extent. For most methods, the removal of monocytes resulted in the deterioration of mDC estimates (i.e., higher RMSE, **Fig. S7**). On the contrary, removing NK cells improved the estimates of T cell subsets. This effect was more evident in the *Hoek* dataset, since the T cells were not divided into similar subpopulations. AutoGeneS was strongly impacted by removing cell types from the reference dataset. Vice versa, Bisque was impacted the least but showed overall low accuracy.

In a related analysis, we assessed the presence of spillover-like effects, i.e., the overestimation of a cell type caused by cell types with similar transcriptional profiles (**Fig. 1C**). We used two real bulk datasets (*Hoek-pure* [44] and *Linsley-pure* [54]) that consist of RNA-seq samples from purified immune-cell populations, which we deconvolved after methods’ training using the *HaoSub* scRNA-seq reference. Scaden showed the lowest cell-type-specific and total spillover in both real and simulated bulk datasets (**Fig. 5A,B, S8, and S9**), challenged only by mDCs in the *Hoek-pure* dataset (**Fig. S9**), where on average 10% were misclassified as monocytes. CIBERSORTx was the only method that could correctly differentiate between Monocytes in mDCs in the *Hoek-pure* dataset, however it incorrectly estimated a large portion (46%) of CD8^+^ T cells towards other cell types (18% to NK cells and 10% to Tregs and CD4^+^ T cells) in the *Linsley-pure* dataset. SCDC and MuSiC also obtained low spillover, mainly restricted to T-cell subtypes Similarly for DWLS, which additionally showed high spillover from B to plasma cells. Only Scaden could correctly differentiate between T cell subtypes in the *Linsley-pure* dataset.

**Fig. 5:**
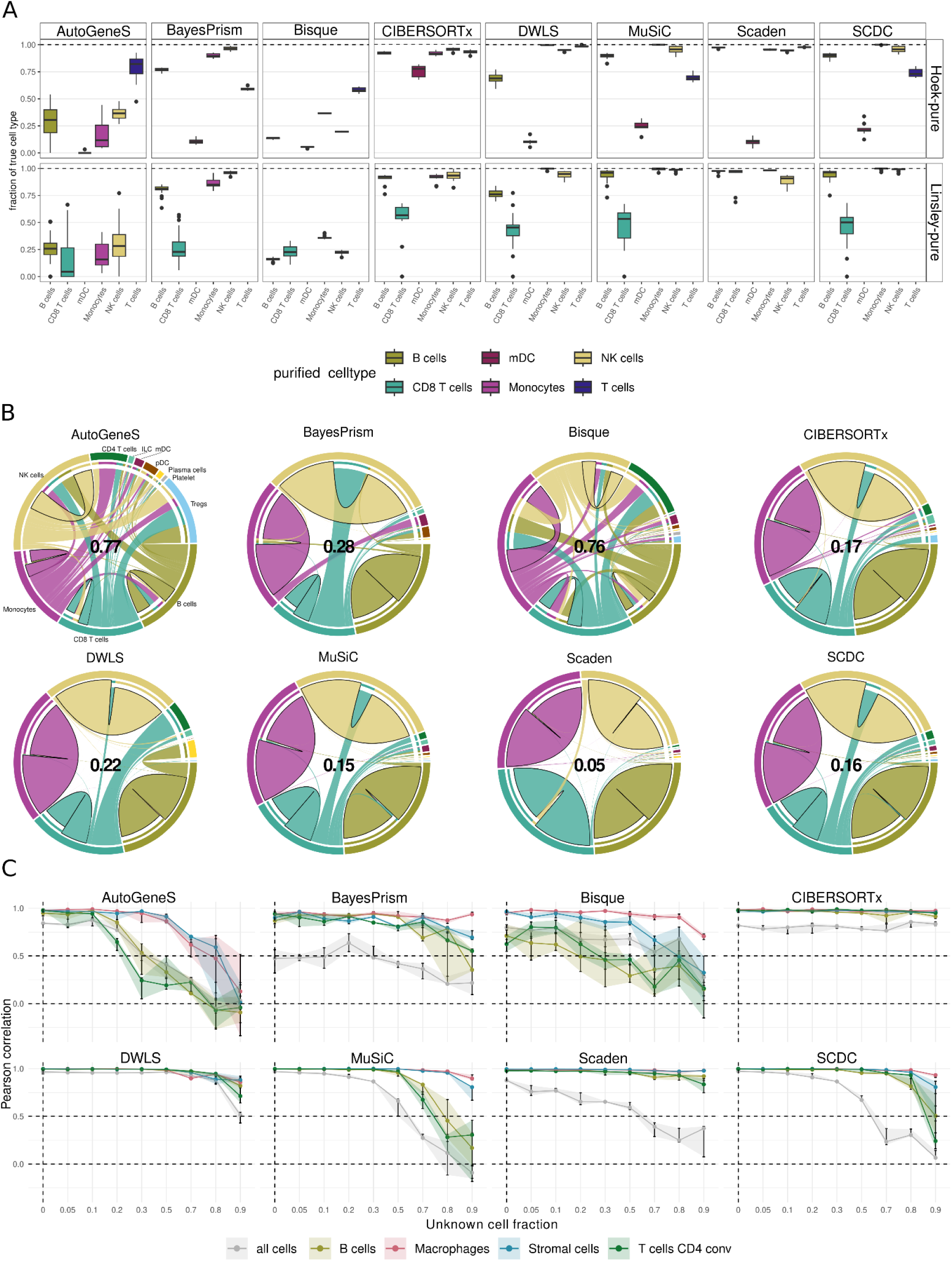
Exploration of different sources for over-estimation include spillover and unknown content in the pseudo-bulk. **(A)** Percentage of correctly predicted cell type abundance in the spillover analysis on the *HaoSub* reference while deconvolving purified bulk RNAseq samples from two datasets (*Hoek-pure* [35] and *Linsley-pure* [45]). The presented values correspond to the sum of the correctly predicted cell fractions for each method. **(B)** Chord diagrams showing the cell type predictions for the spillover analysis for the *Linsley-pure* dataset. The outer circle indicates the different cell types, while the chords represent the mean cell fractions predicted across the various samples. A connection with a bold outer black line, starting and leading to the same segment represents a correct prediction, while a connection leading to a different section indicates the presence of spillover towards this cell type. The values in the center indicate the total fraction of predictions attributed to wrong cell types. **(C)** Performance of the different deconvolution methods in the simulation with unknown tumor content using the *Lambrechts* scRNA-seq reference. Pseudo-bulks (n=50) were simulated using the same single-cell dataset, for each level of unknown tumor fractions. The mean Pearson correlation coefficient was computed for the predictions of each cell type. The shaded areas show the interquartile range as five technical replicates have been generated for each level of tumor content.

To further leverage exactly-matching cell-type labels between single-cell and bulk data, we reproduced the spillover analysis with pseudo-bulk samples. We simulated pseudo-bulk samples with SimBu directly from the *Hao* single-cell reference by aggregating expression profiles of individual cells per cell type, ensuring consistent definitions and enabling a fair evaluation of spillover between closely related populations (Methods). DWLS was the best-performing method, with true fractions above 90% for most cell types (**Fig. S8B**) and only 7% of total incorrect (“spillover”) predictions (**Fig. S8C**); outliers were ILCs (mainly spilling over to CD4^+^ T cells) and T_REG_ cells. MuSiC, BayesPrism, SCDC, and Scaden followed closely by DWLS, with total spillover values from 11% to 16%. They again showed robust estimates for monocytes, NK cells, B cells, mDCs, and CD4^+^/CD8^+^ T cells. However, they were less accurate for T_REG_ cells or plasma cells, which were often classified as CD4^+^ T cells and B cells, respectively. Generally, monocytes, NK, B, and T cells could be estimated with little spillover in real and simulated bulk samples by DWLS, Scaden, SCDC, and MuSiC. When looking into T cell subtypes, only Scaden maintained low spillover, as already confirmed on the real RNA-seq datasets. Bisque failed this test, with large spillover in the myeloid and lymphocyte compartments in both real and simulated settings.

Deconvolution has classically been applied to characterize immune cell infiltration in bulk-tumor RNA-seq samples, which is a key biomarker for clinical decision-making in oncology [1,55]. However, while immune-cell signatures can be easily derived from public data, suitable tumor-cell expression signatures are rarely available. Tumor cells show large variability between and even within patients [56], and tumor-matching scRNA-seq references for method training are rarely available. This leads to a critical and commonly overlooked challenge: bulk RNA-seq datasets often contain unknown or unmodeled cell types, such as tumor cells, which are not represented in the single-cell reference used for methods training. Hence, we evaluated the impact of unknown tumor content in bulk RNA-seq data using pseudo-bulks simulated from the *Lambrechts* lung cancer dataset [3]. We mixed single-cell profiles from B cells, macrophages, CD4^+^ T cells, and stromal cells, and introduced increasing fractions of tumor-cell profiles (Methods). Single-cell data from the same cell types, except tumor cells, was used for training. For each level of tumor content, we evaluated the deconvolution performance using Pearson correlation (**Fig. 5C**). CIBERSORTx showed the most stable results in agreement with its original study [35], with correlations close to 1, regardless of the proportion of unknown content. DWLS and Scaden showed very stable results, with correlations close to 1 up to 80% unknown content for the individual cell types. On global correlation (gray curves), CIBERSORTx was similarly stable, although at a lower performance level (correlation around 0.8). DWLS obtained high correlations here, although it dropped at 80% unknown content. Scaden, MuSiC, and SCDC showed similar patterns, though correlation decreased earlier, at 30-50% unknown content. Overall, global correlation was more impacted than the correlations for the individual cell types, indicating an estimation bias due to the failed quantification of the tumor-cell content and to the resulting overestimation of the quantified cell types. In this analysis, macrophages seemed the easiest cell subpopulation to be quantified for most methods. For BayesPrism and Scaden, stromal cell identification was the most problematic, while AutoGeneS struggled with CD4^+^ T cells.

### Impact of the single-cell reference dataset on deconvolution performance

An important question is to what degree deconvolution performance is affected by the choice of the single-cell RNA-seq reference data set. To systematically investigate this, we first focused on the impact of the reference dataset/study, keeping the single-cell protocol (10x Genomics 3’ v2 or v3) and tissue type (healthy blood cells) fixed (Methods). In addition to the *Hao* dataset, we selected seven additional single-cell datasets satisfying these criteria using the CZ CELLxGENE Discover platform [57] (**Table S2 and Fig. S10**). To ensure a uniform cell-type annotation between all datasets, we integrated them using scANVI [58] and labelled each cluster to a shared cell-type nomenclature leveraging marker-gene expression and our established high-confidence labels of cells from the *Hao* dataset (Methods). Finally, the manually annotated datasets were used to deconvolve the real bulk samples from the *Finotello*, *Hoek, Morandini,* and *Altman* datasets.

The estimation of NK cells, CD4^+^ T cells, and CD8^+^ T cells was generally consistent across methods, with low variability – except for AutoGeneS, which instead showed high variance in its predictions (**Fig. 6A**). For NK cells, for example, the correlation ranged widely, with values of -0.07 ± 0.37 in the *Finotello* dataset, and 0.23 ± 0.42 in the *Hoek* dataset (**Fig. S11-13**). Monocytes and mDCs displayed the highest variability across methods, reference single-cell datasets, and bulk datasets, suggesting that their deconvolution is particularly sensitive to the choice of the single-cell reference. With the exception of mDCs, Scaden consistently showed low variability across cell types, suggesting its robustness to the choice of reference dataset. MuSiC and SCDC showed limited variability, but also an inferior overall deconvolution performance. Interestingly, we observed that the variability across references differed between correlation and RMSE depending on the cell type and method, reflecting distinct aspects of deconvolution robustness. For example, DWLS exhibited low RMSE variability (with the exception of B cells) but high variability of correlation values across cell types, suggesting that while its absolute estimates remain stable, its ability to rank samples correctly is more sensitive to the choice of reference.

**Fig. 6:**
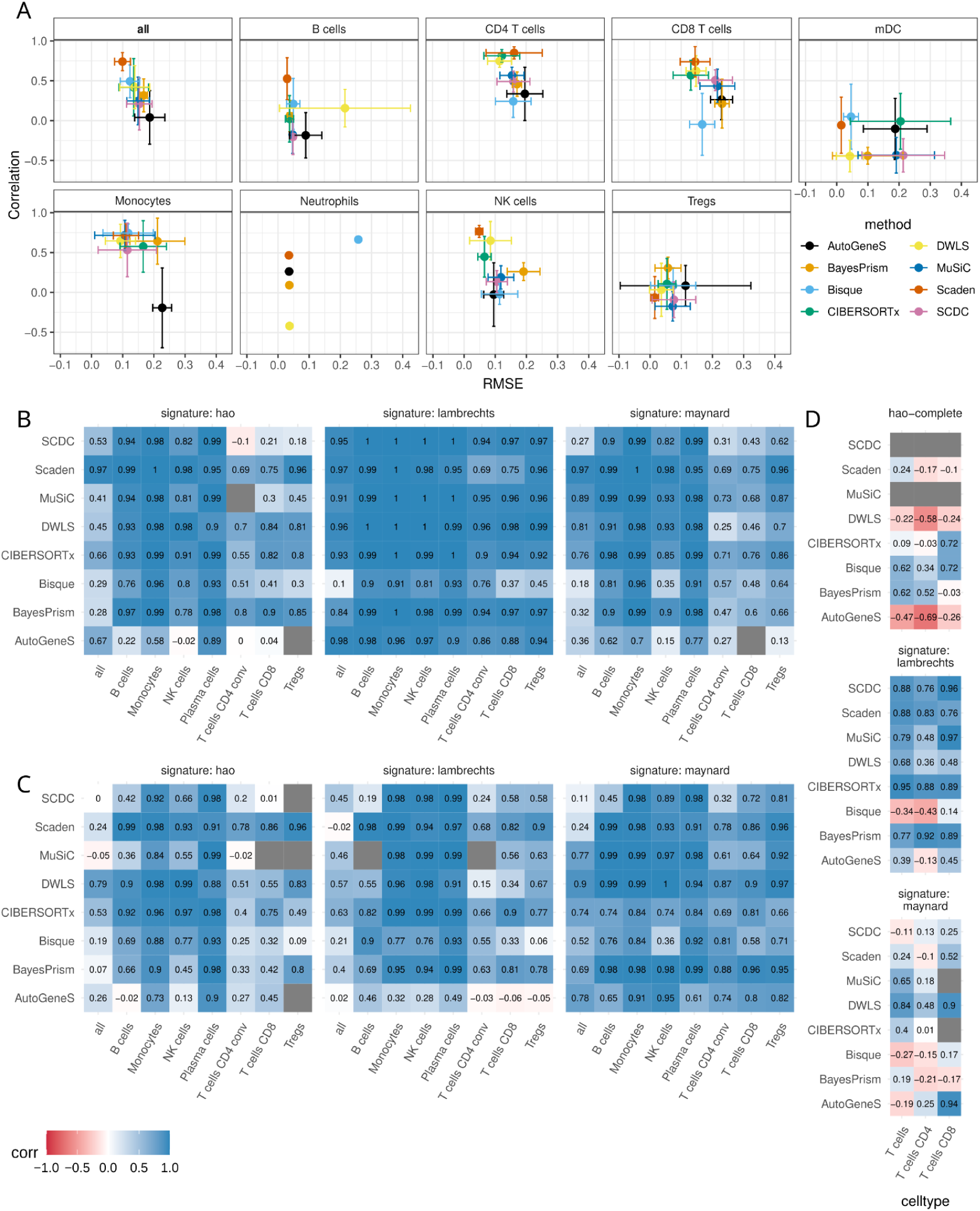
Single-cell reference-based effects on deconvolution performance. (**A**) Mean Pearson correlation and mean RMSE for cell-type predictions across eight different single-cell reference datasets applied to the real *Finotello* bulk dataset. Error bars represent the standard deviations of both metrics for each method in a cell-type, calculated across the eight different references. The first box with ‘all’ shows the correlation and RMSE based on all cell-type fractions combined; Neutrophils could only be detected with one reference dataset (*Tabula Sapiens*). Deconvolution of pseudo-bulks derived from *Lambrechts* **(B)** or *Maynard* **(C),** and the *Vanderbilt* bulk dataset (**D**) with methods trained on different single cell datasets (indicated in headers of each heatmap). The heatmaps show the Pearson correlation coefficient between the simulated ground truth cell fractions and the estimates obtained training the methods with different signatures. Grey tiles correspond to NA correlation values, due to all the estimates for cell type fractions being zero.

In real settings, the performance of scRNA-seq-informed deconvolution might be influenced by two other variables: tissue/disease context and single-cell protocol/technology, which were kept fixed in the previous analysis. To assess the impact of these aspects, we simulated two pseudo-bulk datasets from lung-cancer scRNA-seq datasets generated with different technologies: *Lambrechts* [3] (10x Genomics) and *Maynard* [59] (Smart-seq2). Of note, we leveraged the harmonized cell-type annotations provided by the Lung Cancer Cell Atlas (LuCA) [4] to minimize discrepancies between cell-type labels. To deconvolve these data, we considered either one of these single-cell datasets for training. As tissue-specific references may not always be available, we used the *HaoSub* PBMC dataset as a reference (10x, PBMC). For both pseudo-bulk simulation and methods training, only the cell types in common across all three single-cell datasets were considered. As expected, the most accurate results were obtained with matching pseudo-bulk and single-cell reference (**Fig. 6B, C**). When using training data from a different single-cell technology, the deconvolution performance deteriorated. The worst results were obtained when the training data came from a different tissue context (i.e., *HaoSub* PBMC data), with some T-cell subsets being undetected in the pseudo-bulks. B cells, monocytes, plasma cells, and, in some settings, NK cells seemed less affected by the technology and tissue context. Similarly, some methods (Scaden, DWLS, and CIBERSORTx) proved to be more robust to the impact of technology.

For signature-based (rather than model-based) methods, some determinants of deconvolution performance can be investigated in the characteristics of the derived signature matrix. Although signature assessment (only possible for half of the methods considered in this benchmarking) is beyond the scope of this study, we show an example of how our interactive web app *deconvExplorer* (https://www.daisybio.ls.tum.de/deconvexplorer/) can be used to compare the signatures derived by DWLS from the two lung-cancer scRNA-seq datasets (**Fig. S14**, Methods). Both signatures contained similar numbers of genes (*Lambrechts*: 1,112, *Maynard*: 1,331), though only 359 genes were in common. The lower condition number of the Lambrechts signature (357.52) compared to the Maynard (502.28) indicates a higher discriminative power, in line with the obtained deconvolution results (**Fig. 6B-D**). Finally, visual inspection of the signatures represented as z-scored heatmaps shows clearly the unspecificity of various genes across T cell subtypes, additionally quantified by a lower gene-wise Gini index. The *deconvExplorer* web app can aid in the manual curation of expression signatures and removal of genes with low cell-type specificity, which is an effective strategy to reduce the spillover effect and increase the deconvolution performance [8].

In addition to the pseudo-bulk analysis, we used all three single-cell reference datasets (this time using all available cell types) to deconvolve bulk RNA-seq data from lung tumors for which CD4^+^ and CD8^+^ T cell estimates were available from IHC (*Vanderbilt* dataset [43], **Fig. 6D**). Most methods showed the poorest results using the *HaoSub* PBMC data set, confirming the importance of having training data from a matching tissue and disease context. The impact of the single-cell technology was instead less clear in this assessment. Most methods performed better on the *Lambrechts* dataset, with the exception of Bisque and AutoGeneS. This real-life application also shows that all methods struggled to robustly estimate tumor-infiltrating lymphocytes, which are transcriptionally similar and rare compared to the overall sample composition (CD4^+^ T cells: 0.03±0.07%; CD8^+^ T cells: 0.01±0.06%).

## Discussion

Second-generation deconvolution is a powerful technique to investigate the complexity of tissues, coupling the high resolution of single-cell data [4,60–63] with the cost-effectiveness of bulk RNA-seq [18]. In principle, second-generation deconvolution methods can be applied to any cell type, tissue, and organism, but their inherent flexibility challenges their validation and the understanding of the determinants of deconvolution performance. In this study, we benchmarked a selection of second-generation deconvolution methods. Instead of focusing on previously-assessed aspects like data normalization [20], we addressed a set of open challenges in second-generation deconvolution [25,26], which guided us in the rational design of a compendium of real and simulated bulk RNA-seq datasets across different tissues and organisms (**Fig. 1C**). Bulk datasets with reliable cell composition information are scarce and, due to different biases introduced by experimental protocols, the measured FACS proportions might not reflect those present in the bulk RNA-seq [25,26,64], especially for fine-grained cell types [65]. However, they still serve as the current gold standard in the field. On the other hand, some of our simulated datasets might not be representative of real-life situations, but they nonetheless help to disentangle the impact of various sources of variability and understand methods’ behavior under a variety of controlled scenarios, which would be unfeasible to reproduce experimentally. We deliberately disregarded “validation” datasets whose ground-truth cell-type fractions were derived from scRNA-seq or from the sequencing of admixed RNAs (rather than cells). Data from scRNA-seq studies currently cannot be regarded as a gold standard due to biases in cell capture efficiency [3,6]. The underrepresentation of neutrophils in current single-cell studies serves as a prominent example [4], but other cell types like adipocytes are affected as well [5].

In our assessment based on real bulk RNA-seq data, Scaden and DWLS consistently performed best. Nevertheless, they still returned inaccurate estimates for certain datasets and cell types, which would be hard to spot in real-world investigations, where the true cell fractions are not known a priori. For this reason, we would recommend running and comparing a few top-performing methods to spot early on possible inconsistencies in the predictions. The *omnideconv* R package provides a convenient framework to support such comparative analyses. In the absence of a guiding ground truth, users should pay attention to other “red flags” like extremely high (i.e., non-physiologic) cell fractions, systematically absent cell types, or unstable results that depend heavily on the scRNA-seq reference used for training. Notably, we observed large discrepancies between CIBERSORTx runs executed via the Docker container (as supported by *omnideconv*) and those performed through the web application. However, we were unable to systematically test the latter due to upload limitations on the size of single-cell reference data and the impracticality of running large-scale analyses through the web interface.

Another feature of the input scRNA-seq data that influences method performance is the resolution of cell-type annotations. A goal of cell type deconvolution methods, especially in biomedical research, is to discern not just broad cell types but also to resolve detailed sub-types or functional states. In agreement with a previous study [42], granular cell types, especially T-cell subsets, were more difficult to quantify for most methods. DWLS consistently provided the most accurate results when tested on the deconvolution of fine-grained cell types from different tissues. Of note, in the deconvolution of coarse cell types, our analysis across different tissues showed that methods like AutoGeneS and CIBERSORTx benefit from training on fine-grained-annotated scRNA-seq data and a posteriori aggregation of the estimates for coarser cell types, in agreement with previous studies [31,34].

Besides prediction accuracy, computational scalability is a key factor in selecting the preferred method for real-world applications. Our results highlight substantial computational differences between methods. DWLS and Scaden were among the most demanding, while AutoGeneS, SCDC, and MuSiC were faster and more memory efficient. Since signature creation was the main computational bottleneck (including DWLS and Scaden), reusing signatures across similar datasets offers a practical way to reduce runtime, which is supported by *omnideconv*.

The reduction of the reference dataset size is a promising way for extracting precious information for deconvolution analysis from large single-cell atlases, but it can be achieved with different approaches, which can have a different impact on deconvolution performance. For instance, single cells can be sampled evenly or maintain the original proportions across cell types. Other strategies that can impact deconvolution results in a method-specific manner [66] are the use of “metacell” aggregation [67,68] or the selection of highly representative cells from integrated datasets [69]. A thorough evaluation of these alternative strategies would be required to develop robust, method-specific guidelines for single-cell reference reduction – an effort that falls beyond the scope of this work and would merit a dedicated study.

In this study, we investigated three major challenges that can result in biased deconvolution results: cell-type-specific mRNA bias, transcriptional similarity of closely-related cell types, and presence of unknown cellular content. Cell-type-specific mRNA bias deriving from different cell sizes and transcriptional activities can result in systematic estimation bias of cell fractions and, thus, must be correctly handled by deconvolution methods [1]. Leveraging our pseudo-bulk simulator SimBu [27], we showed that only DWLS, MuSiC, Scaden, and SCDC robustly corrected for mRNA bias. Methods that are not robustly correcting this bias could be possibly coupled with suitable normalization strategies [70,71].

A major challenge for deconvolution methods is an incomplete single-cell reference. scRNA-seq data are affected by cell-dissociation biases that distort the true cellular composition [3], with potential loss of specific cell types such as neutrophils [4,72]. In our evaluation, the absence of a specific cell type severely impacted the estimates, especially in the presence of transcriptionally similar cell types –a finding recapitulated by our “spillover” analysis. Moreover, the spillover analysis, quantifying the proportion of “background predictions” assigned to incorrect cell types, highlighted that methods like Bisque are clearly optimized for scenarios where bulk and single-cell data are matched, or at least where cell-type proportions fall within a similar range, as also observed by Huuki-Myers et al. [53]. Since such matched datasets are relatively rare, this constrains the utility of this approach in typical deconvolution scenarios that involve data from independent sources and, especially, the quantification of disease-induced cellular shifts.

The robustness to unknown cellular content is key in ensuring between-sample comparability, especially in tumor-bulk deconvolution. In the absence of appropriate tumor signatures, first-generation methods for immune-cell type deconvolution, such as quanTIseq [43] and EPIC [14], enable the quantification of the percentage of “uncharacterized cells”, which in bulk-tumor deconvolution can be regarded as a proxy for tumor content [43]. None of the considered second-generation deconvolution methods offer this feature, as they all normalize cell type fractions to sum up to 100% of the sample. Nevertheless, our results indicate that tools like CIBERSORTx, DWLS, and Scaden can provide estimates that are positively correlated with the true cell fractions even when the unknown tumor content reaches 80-90% of the sample composition. We point out, however, that the estimates, even when positively correlated with the ground truth, have to be considered as systematically inflated due to the failed quantification of the unknown content. This highlights the pressing need for novel deconvolution approaches that explicitly model and correct for unknown cellular content in order to ensure accurate and interpretable quantification across diverse tissue samples.

For commonly-studied tissues such as healthy blood, multiple publicly available scRNA-seq references exist [57], but it is unclear how robust deconvolution methods are to between-study technical and biological variation. Using eight healthy blood references for methods training, we found that overall method performance varied across references, with monocytes being the most impacted cell type. Instead, methods that struggled to estimate certain cell types did so consistently, regardless of the reference used. Nevertheless, Scaden and DWLS showed the lowest variation across the tested references, bulk datasets, and cell types. Interestingly, performance variability was more pronounced for correlation than for RMSE, suggesting that certain methods are prone to systematic estimation for certain cell types, regardless of the reference used. Of note, the more stable results obtained for large bulk datasets like *Morandini* (n=113) and *Altman* (n=332) could be partly explained by a more limited impact of outlier samples compared to smaller datasets such as *Finotello* (n=9) and *Hoek* (n=9). In this study, we leveraged the shared cell-type annotations curated after single-cell data annotation with scANVI [58]. Still, every single-cell reference has been used independently from the others for methods training and downstream deconvolution. As (multi-study) single-cell projects and the integration into single-cell atlases becomes more popular, this raises new opportunities for learning robust signatures for cell-type deconvolution. However, there are also open questions to be investigated, e.g., how to address batch effects between studies or whether normalized data and embeddings (which are common for single-cell atlases) are suitable for this task.

Finally, our findings suggest that the tissue/disease context of the single-cell data used for method training can have a major impact on deconvolution results, while the effects of the sequencing technology were more variable across methods and datasets. When deconvolving lung cancer pseudo-bulks with PBMC signatures, some cell types were often undetected, emphasizing tissue-specific differences in the transcriptomic profiles of various cell types [73] and the need for a biologically similar background of the single-cell and bulk RNA-seq data. Our results also suggest that tag-based sequencing data (e.g.,10x) represents the most appropriate reference, though certain methods produce more accurate results when trained with full-transcript data (e.g., Smart-seq2). We speculate that this may be related to the data type used for method development and validation. While we focused here on the characteristics of the scRNA-seq reference used for training, the experimental protocols used for the generation of the bulk RNA-seq dataset to be deconvolved can also affect the final results, as investigated in a previous study [53].

## Conclusions

In our assessment, DWLS and Scaden showed the most robust deconvolution performance across multiple scenarios; especially noteworthy is the amount of unknown content and spillover they can handle. Nevertheless, they also showed decreased accuracy in some settings, and they both required the largest computational resources across the tested methods. As method performance is strongly context-specific, we do not provide a definitive ranking or recommendation.

To ensure a fair comparison and avoid biasing results toward any particular method, we deliberately refrained from parameter tuning, relying instead on developer-recommended or default settings. While this approach supports reproducibility, future studies could explore how parameter optimization and method-specific features influence performance, facilitated by the benchmarking tools we provide through https://omnideconv.org/. For instance, some methods could possibly improve their deconvolution performance through more tailored data and deconvolution approaches, like multi-level cell-type annotation (BayesPrism), matching bulk and single-cell RNA-seq data (Bisque), multiple scRNA-seq datasets (Scaden and SCDC). Another possible improvement would involve the pre-selection of marker genes for certain cell types to aid the methods in creating a more robust and representative signature [53] or, for methods like CIBERSORTx, the provision of curated/optimized gene signature matrices [74]. All methods included in *omnideconv* can be easily used to their full potential and, together with our additional novel resources, we envision researchers having better possibilities to thoroughly characterize and, especially, optimize deconvolution results, extending this powerful technique to an increasing panel of applications and domains.

## Methods

### Bulk RNA-seq data accession and processing

Raw FASTQ files were retrieved from the Gene Expression Omnibus (GEO) using the accession codes GSE107572 (*Finotello [43]* dataset, n=9), GSE64655 (*Hoek [44]* dataset, n=8 PBMC samples and *Hoek-pure* with n=48 purified cell populations), GSE115823 (*Altman* dataset, n=322), GSE193142 (*Morandini* dataset, n=159) and GSE60424 (*Linsley*, n=114 purified cell populations) as well as ArrayExpress using the accession codes E-MTAB-9271 (*Petitprez [49]* dataset, n=14) and E-MTAB-6458 (*Chen[48]* dataset, n=12). The datasets were processed in the same way, using the nf-core RNA-seq framework [75] (v3.10.1) and the human reference genomes GRCh38 with genome annotation v41, and murine genome GRCm39 with genome annotations vM30, from the Genome Reference Consortium. Briefly, FASTQ files were first aligned to the reference genome with STAR [76] (v2.7.10), and then summarized with Salmon [77] (v1.9.0) to obtain both read counts and transcripts per million (TPMs). The *Vanderbilt* dataset was provided by the Vanderbilt Institute for Infection, Immunology, and Inflammation (USA) [43]. The flow cytometry estimates for the various cell types were obtained from the corresponding authors. More details on the bulk datasets are available in **Table S1**.

All bulk datasets that have been generated and are used in this work are deposited in *deconvData* as a convenient resource to use for further deconvolution tasks.

### Single-cell RNA-seq data accession and processing

The count matrix and the cell type annotations for the *Hao [47]* and the *Wu [50]* single cell datasets were retrieved from GEO, with accession numbers GSE164378 and GSE176078, respectively. Both the *Lambrechts [3]* (number of cells = 64,135) and the *Maynard [59]* datasets were retrieved from the LuCA atlas (https://luca.icbi.at/, V. 2022.10.20) [4], and the cells classified as doublets (‘doublet_status’) were removed. The Tabula Muris (*TM*) spleen dataset was retrieved from the atlas website (https://tabula-muris.ds.czbiohub.org/) [60]. The *Allen* brain dataset [51] was downloaded from the Allen Brain Map website (https://portal.brain-map.org/atlases-and-data/rnaseq). Additionally, we accessed a set of scRNA-seq datasets from healthy adult blood samples that have been processed with the 10x 3’ protocols using CZ CELLxGENE census [57] (census_version="2025-01-30"). Three datasets of this list were further excluded, due to having samples from pregnancy or only two biological replicates. Further, we excluded datasets that have fewer than 5,000 or more than 200,000 cells. The above-mentioned *Hao* dataset was part of this list of remaining datasets; however, we continued with the expression values from GEO, as this dataset has additional information on surface protein expression, based on the CITE-seq assay.

The single-cell datasets were preprocessed with Seurat [78] and scanpy [79]. Quality control was performed according to the current best practices [80], considering the number of total counts, the number of expressed features (genes), and the fraction of mitochondrial genes per cell. We considered the MAD (Median Absolute Deviation), computed as 𝑀𝐴𝐷 = 𝑚𝑒𝑑𝑖𝑎𝑛(|𝑋 − 𝑚𝑒𝑑𝑖𝑎𝑛(𝑋)|), with 𝑋 being the respective metric of an observation. Cells that have 𝑋 < 𝑛 * |𝑀𝐴𝐷|, with 𝑛 = 5 for the number of total counts and the number of expressed genes, or for the fraction of mitochondrial RNA were classified as outliers.

Single cell datasets were CPM-normalized through Seurat. For the *Maynard* dataset, transcript-length-normalized values were retrieved from the LuCA atlas (‘counts_length_scaled’) and were then CPM-normalized. To create a unified cell-type annotation across all eight described healthy human blood datasets (including *Hao*), we first manually unified the original cell-type annotations into the following labels: T and NK cells, Myeloid cells, Neutrophils, B cells, red blood cells, Platelets, pDCs, HSCs, ILC, or Basophils. We used scANVI [58] to integrate the datasets over the donor ID as batch variable and our manual annotations, using the nf-core scdownstream pipeline (https://nf-co.re/scdownstream/dev/) (**Fig. S10**), which also prepared Leiden clusters in different resolutions. Subsequently, we removed all non-expressed genes. More details on the scRNA-seq datasets are available in **Table S2**.

### Single-cell RNA-seq data annotation

For the *Maynard* dataset, the cell type annotation considered was the ‘cell_type_tumor’ column. We removed all cell types that were labeled as ‘dividing’ and ‘transitional’. For the *Lambrechts* and *Maynard* dataset, the medium and fine cell type annotations considered were the ‘cell_type_tumor’ and ‘ann_coarse’ columns, while the coarse level was obtained aggregating the dendritic cells, macrophages /monocytes and T/NK cells populations. The coarse and fine cell type annotations for the *Wu* dataset were obtained from the metadata provided with the cell counts (‘celltype_major’ and ‘celltype_minor’ fields, respectively). The T cell subpopulation in the *TM* dataset was manually annotated to identify the CD4^+^, CD8^+^ and Tregs subpopulation. A detailed table of cell-type labels can be found in **Table S3**. The original cell-type annotations of the *Hao* dataset where refined using the gene and surface protein expression and used as basis for the annotation of other blood scRNA-seq datasets: we manually inspected each Leiden cluster (resolution=4) and annotated them manually based on marker gene expression and the established *Hao* ground-truth labels. We did not annotate some ambiguous clusters, where no clear label could be established, and kept these cells as ‘unassigned’. The integrated datasets along with their annotations are available as interactive UMAP via CELLxGENE at https://www.daisybio.ls.tum.de/omnideconv/app/blood.

To use these datasets as references for deconvolution, we removed cells that were annotated as HSCs or were left ‘unassigned’. For the *Hao* dataset specifically, we also removed cells that in the end had ambiguous annotations between our manual ground-truth labels and the labels based on cluster annotation in the integrated blood scRNA-seq datasets. As the final *Hao* dataset was too large to be used as reference for every deconvolution method (147,391 cells), we created a subset (*HaoSub,* 14,744 cells) that contains 10% of each cell type, stopping the subsampling at cell-types with less than 20 cells. We wanted to ensure that the remaining blood scRNA-seq datasets would be similar in size and also subsampled them to 15,000 cells, while keeping cell-type proportions matched to the respective full dataset and ensuring that cell-types with less than 20 cells were not subjected to the downsampling process.

All scRNA-seq datasets that have been generated and are used in this work are deposited in *deconvData* as a convenient resource to be used for further deconvolution tasks.

### Benchmarking pipeline *deconvBench*

We implemented a Nextflow pipeline called *deconvBench* that helped us to perform all of the simulation scenarios described above in an efficient and reproducible way: https://github.com/omnideconv/deconvBench. This pipeline was executed on a cloud computing cluster provided by the German Network for Bioinformatics Infrastructure (de.NBI).

### Performance metrics

We considered two primary metrics for the performance of the benchmarked methods: Pearson correlation and root mean square error (RMSE). Both were calculated by comparing the available ground truth cell-type fraction against estimates from the methods under evaluation. Metrics were calculated across all cell-type fractions and samples (*global correlation*) and separately for each individual cell type, facilitating both global and cell-type-specific assessments.

For each sample *s* with vectors *x* (ground truth cell-type fractions) and *y* (estimated fractions by a deconvolution method) of the same length *C* (number of cell types), we calculated sample-wise RMSE as 𝑅𝑀𝑆𝐸 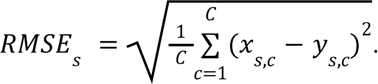. Similarly, we calculated RMSE for each cell-type *c* across all available samples *S* as 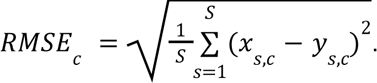.

RMSE together with correlation allowed us to quantify the *estimation bias* of methods, i.e.,the systematic deviation of predictions from the ground truth in a positive (over-estimation) or negative (under-estimation) direction. We additionally calculate two other metrics in the *deconvBench* pipeline, mean absolute error (MAE) 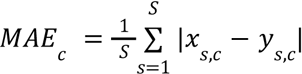 and mean absolute percentage error (MAPE), as 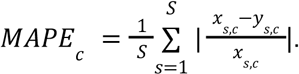

### Simulation of pseudo-bulk RNA-seq datasets

Pseudo-bulk RNA-seq datasets were simulated from scRNA-seq data to replicate realistic bulk RNA-seq profiles with known cell-type compositions. Using the *Hao* scRNA-seq dataset, we prepared pseudo-bulk datasets, in which we mirrored sample-specific features of real bulk datasets (*Finotello*, *Hoek*, *Morandini,* and *Altman*) with matching flow cytometry or CBC-derived cell fractions. Each real bulk sample served as a template for 10 pseudo-bulk technical replicates (5 in the case of *Morandini* and *Altman* due to their already larger sample size). Similarly, we used two mouse datasets (*Chen* and *Petitprez*) for which the corresponding cell fractions were available, along with the Tabula Muris (*TM*) spleen scRNA-seq reference. The simulations were carried out with the R package SimBu [27] (version 1.8) with the following settings:

Each pseudo-bulk sample sampled 10,000 cells (with replacement) from the scRNA-seq datasets to have the same cell type composition as the bulk sample *n* (derived from FACS data). Only cell types present in both the real bulk and scRNA-seq datasets were utilized for simulation. The pseudo-bulks were simulated to have the same sequencing depth as the bulk sample *n*. Additionally, we scaled the expression of genes in each cell by scaling factors derived from the total number of expressed genes per cell in the scRNA-seq reference dataset (**Fig. S1**). This approach has been established as a ‘silver standard’ for accounting for mRNA content bias in cells [27].

As there are missing values in the FACS annotation in four samples of the *Petitprez* dataset, we left out these samples in the described setup. The *Hoek* bulk dataset only includes FACS annotations for T cells, but the *Hao* reference differentiates three subtypes (CD4+ T cells, CD8+ T cells, and T regulatory cells). Therefore, fractions estimated for these T cell subsets were aggregated for comparison purposes. For the Hoek-based pseudo-bulks (called *HoekSim*), we sampled from all three T cell subtypes in *Hao* during simulation. The *Morandini* bulk dataset contains FACS annotations for different levels of cell types, such as Lymphocytes and their subtypes. Similarly to the *HoekSim* pseudo-bulk, we only selected these subtypes from the Hao reference to be used during the simulation of *MorandiniSim*. For the *AltmanSim* pseudo-bulks, we sampled from CD4 T cells, CD8 T cells, Tregs, B cells, and ILCs to form the Lymphocyte population and from Monocytes, mDCs, and pDCs for the Monocyte population. Both the *MorandiniSim* and *AltmanSim* are based on more extensive real bulk datasets; therefore, we only created 5 pseudo-bulk technical replicates instead of 10.

### Simulation and deconvolution of pseudo-bulk RNA-seq datasets to study mRNA bias

To explicitly evaluate mRNA content bias effects, we used the previously described blood-derived scRNA-seq reference datasets (**Table S2**) to simulate a set of pseudo-bulk datasets with SimBu. To confirm that our results would in the end also generalize to other tissues, we executed the same procedure on the *Allen* brain scRNA-seq dataset. From each reference, we sampled 10,000 cells in random proportions from the present cell types, creating 50 pseudo-bulk samples, fixing the simulated sequencing depth per sample to 10,000,000. We executed this simulation setup twice, once with the number of expressed genes as cell-type-specific scaling factor (as described also above) and then again without any mRNA bias scaling (i.e., the *scaling_factor* factor parameter set to NONE). Additionally, to ensure comparability between samples, we used the same fixed seed for each sample with and without mRNA bias, such that the same cells are sampled for both samples, and the only difference is due to mRNA bias modelling. In the end, we prepared a compendium of 800 samples to test the deconvolution methods rigorously on different tissues and cell types.

The pseudo-bulks with and without mRNA bias scaling were then deconvolved using the same scRNA-seq reference dataset that was used to create the samples. This would ensure that method performances generally remain on a high level, and we could focus only on performance differences due to mRNA bias. Finally, we compared the estimated cell-type fractions from the simulations with and without mRNA bias with the simulated ground truth to analyse how well each method can account for mRNA bias. Whenever the performance would decrease on the simulations without bias, we would consider the method to correctly account for mRNA bias for a specific cell type (and vice-versa).

### Measuring computational efficiency

To benchmark methods resource requirements based on a change in scRNA-seq reference size, we subsampled the complete *Hao* dataset to different subsets of cells (1000, 3000, 5000, 10000, 25000, 50000, 100000). These subsets were used as input for the deconvolution methods to establish how much memory usage and runtime would increase when more cells are present in the reference. Additionally, we generated five different sets of pseudo-bulk datasets, consisting of different numbers of samples (5, 25, 50, 100, 200) to also monitor changes in resource usage when more samples have to be deconvolved.

Using our deconvBench pipeline, written in Nextflow (version 22.10.1), we were able to measure memory consumption with Nextflows ‘--with-trace’ parameter. It stores - among others - information on consumed memory in the ‘peak_vmem’ column. We calculated the runtime of methods by the difference in the end and start time of the core *omnideconv* functions ‘build_model’ and ‘deconvolute’. In the case of omnideconv deconvolution methods reliant on Python, an overhead may arise as they are invoked through the *reticulate* interface rather than directly. Furthermore, for these methods, R data types must be first converted to formats compatible with Python, adding an additional computational step.

Each method was given a single CPU to perform calculations, we did not use potential parallelization features of the methods. The methods were then executed on an HPC cluster, using only computing nodes with the same hardware and software setup: the nodes use an Intel Xeon Gold 6148 processor with 2.4 GHz, have a maximum accessible RAM of 384Gb and all run Ubuntu 22.04.1 as operating system. All methods were executed inside an apptainer [81,82] (version 1.3.3) container, with the exception of CIBERSORTx, which instead was executed inside a Docker container, though with the exact same versions of dependencies installed. The container we used is available at *deconvData* and *deconvBench*.

### Simulation and deconvolution of pseudo-bulk RNA-seq datasets for spillover analysis

The pseudo-bulk datasets for the spillover analysis were simulated using the *Hao* scRNA-seq dataset. The cell types considered were: B cells, mDC, pDC, CD4^+^ and CD8^+^ T cells, T_reg_ cells, monocytes, NK cells, ILC, platelets, and plasma cells. For each cell type, a pseudo-bulk with 50 replicates was simulated with SimBu using the “pure” scenario and 1000 cells, meaning only cells from this one cell type were sampled. In cases where less than 1000 cells per cell type are available, SimBu is designed to allow the sampling of the same cells multiple times. These pseudo-bulks were then deconvolved with the *HaoSub* reference.

We quantified the amount of spillover as the sum of cell-type estimates that correspond to the pure, simulated cell type (giving a cell-type specific percentage value, ideally 100%) and the sum of cell-type estimates that correspond to all other predicted cell types (giving a method specific percentage value, ideally 0%).

### Simulation of pseudo-bulk RNA-seq datasets with unknown content

The pseudo-bulk datasets for the unknown content analysis were simulated using the *Lambrechts* scRNA-seq dataset. The cell types considered were: B cells, CD4+ T cells, stromal cells, and macrophages, in addition to the tumor cells. The pseudo-bulks were simulated with SimBu using the “weighted” scenario, and increasing concentration of tumor cells: 0%, 5%, 10%, 20%, 30%, 50%, 70%, 80%, 90%. The remaining four cell types were sampled with random fractions. Five replicates with ten pseudo-bulk samples each, were simulated for each tumor concentration, resulting in 9*50=450 pseudo-bulk samples for the deconvolution, the methods were trained with the *Lambrechts* dataset with the four above-mentioned cell types, excluding the tumor cells, downsampled to 500 cells per cell type.

### Simulation of pseudo-bulk RNA-seq datasets for the impact of cell type resolution analysis

The pseudo-bulk datasets to study the impact of cell type annotation resolution were simulated using the *Lambrechts*, *Wu*, and *Allen* scRNA-seq datasets. For the *Lambrechts* dataset, the cell types considered at the “coarse” resolution were: B cells, dendritic cells, monocytes/macrophages, monocytes, and T/NK cells. Except for the B cells, the other cell types possess two additional levels of annotation, “normal” and “fine” (**Table S3**).

Five replicates of ten pseudo-bulk samples each were simulated with SimBu using the ‘mirror_db’ scenario (resembling the cell-type proportions from the scRNA-seq dataset in the pseudo-bulk dataset), considering the “fine” level of annotations. The simulated ground truth estimates of all subtypes of a certain cell type could then be summed up to get the corresponding fractions of this cell type, respectively, for the “normal” and “coarse” annotation levels.

The same was done for the *Wu* dataset, considering (at the coarse resolution) B cells, Cancer-associated fibroblasts (CAFs), Myeloid and T cells (**Table S3**).

For the *Allen* dataset, all cell types were considered for the simulation, and were then grouped to get coarse cell fractions for astrocytes, excitatory and inhibitory neurons, microglia, endothelial cells, oligodendrocytes, and precursors (OPCs). For the visualization of the results, only inhibitory and excitatory neurons were considered, but their fractions refer to the mixture of the whole set of cell types.

This simulation scenario, therefore, provided us with ground truth cell fractions for each pseudo-bulk sample, at three (*Lambrechts*) and two (*Wu, Allen*) resolution levels. For the training of the methods, we only considered the cell types that were also used for simulations and downsampled each cell type to 500 cells, considering the fine cell type annotation. The training datasets were therefore composed of the same count matrix and the different cell type annotations. The pseudo-bulks were then subsequently deconvolved, providing a set of three (*Lambrechts*) or two (*Wu*) cell type fraction estimates.

Finally, we could compare the estimates for cell types that were calculated on each resolution to the corresponding ground truth resolution.

### Simulation of pseudo-bulk RNA-seq datasets for the impact of single-cell technology and tissue context analysis

The pseudo-bulks for the impact of cell technology and tissue context analysis were simulated using the *Lambrechts*, *Maynard,* and *Hao* datasets, considering only the common cell types between the three (B cells, NK cells, CD4^+^ T cells, CD8^+^ T cells, Tregs, Monocytes, and Plasma cells). This was done in order to minimize the variability between the pseudo-bulks and to isolate the effect of the tissue/technology of the dataset. Five replicates of ten pseudo-bulks each were simulated starting from the three single-cell datasets. The same datasets, subsampled at 500 cells per cell type, were then used to train the methods for the deconvolution of the pseudo-bulks. Each pseudo-bulk was then deconvolved with both its corresponding reference and the references from the other two datasets. The Vanderbilt samples, on the other hand, were deconvolved training the methods with the datasets, including every cell type (i.e., also those not used in the pseudo-bulk simulation), subsampled at 500 cells per cell type.

### DeconvExplorer

*deconvExplorer* is an R Shiny web app that aims to provide easy access to analyze deconvolution signatures and results. Signatures can be explored using interactive heatmaps and compared to signatures of other references or methods via upset plots or signature-specific scores such as entropy, condition number, and number of included genes. Finally, deconvolution results can be compared to results of other methods or to user-provided ground-truth data by correlation and RMSE metrics and fitting visualizations. The mentioned scores are calculated as such:

We calculate gene-wise entropy for each gene using the function *entropySpecificity* from the BioQC package [83]. It quantifies the amount of specificity a gene has for a certain cell type, therefore giving information on its potential as a cell-type marker. Genes with a high entropy value (close to 1) will therefore be the best markers, while genes with low entropy values (close to 0) will show a uniform expression level across cell types.

Similarly, the Gini Index is calculated using the same package and its *gini* function. It measures the dispersion in a list of values, where a value of 0 will appear in case that all values are exactly the same. It is maximized (gini_index=1) when the values are maximally different.

The condition number of the signature matrix is calculated using the *kappa* function in base R. It quantifies the stability of the signature matrix towards changes and errors in the input data and is defined as the product of the 2-norm of a matrix *M* and the 2-norm of its inverse:

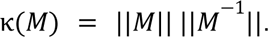

A low condition number (close to 1) suggests that the signature matrix is stable towards changes in the bulk RNA-seq input data. This means that, if the bulk RNA-seq is perturbed by errors, the estimated cell-type fractions will not dramatically differ.

### Details on method settings

The following section covers the versions, parameter settings (if other than default), and data input normalizations we used for each benchmarked method. For some methods, we forked the original repository and had to manually introduce changes to the source code or dependency versions, so that we can reliably install them via *omnideconv*. None of these changes will however influence the results a method produces. Wherever possible, we split the proposed workflow of a method into signature building and deconvolution. Namely, in this benchmark this includes CIBERSORTx, DWLS, SCDC and Scaden.

### AutoGeneS

We cloned the Python package AutoGeneS version 1.0.4 from https://github.com/theislab/AutoGeneS and installed it via the omnideconv package. The method offers an option to only keep highly variable genes for subsequent signature building and deconvolution, however, in order to have the same set of input genes for all methods, we decided not to use this option. Default parameters in omnideconv were used for signature building and deconvolution, only the ‘max_iter’ parameter was fixed to 1,000,000 in order to limit the number of iterations in case no earlier convergence could be found. Also, we set the ‘ngen’ parameter to 5,000, which fixes the number of generations that the algorithm will run. We CPM-normalized scRNA-seq data and TPM-normalized bulk RNA-seq samples, as indicated in the original publication of the method.

AutoGeneS would be able to follow *omnideconv’s* two-step design of separating signature building and deconvolution. However, by far the most amount of runtime is spent for saving the signature as a pickle file. Therefore, we opted to implement it as a one-step method and have the signature creation step as an optional separate step. For the runtime analysis, we benchmarked the one-step version of AutoGeneS in omnideconv, to not artificially increase runtime due to unnecessary IO steps.

### BayesPrism

We forked the R package BayesPrism v2.0 from https://github.com/Danko-Lab/BayesPrism and installed it via the omnideconv package. BayesPrism would offer an option to distinguish between cell types and more detailed cell states in a cellular annotation. As it is the only method in this benchmark with this option, we chose to only use their predictions on cell types in order to keep results comparable. BayesPrism can apply a final round of Gibbs sampling to update the initial cell-type fractions. As recommended by the authors, we did not use this setting in the case that scRNA-seq and bulk RNA-seq originate from similar assays. This is the case when we created pseudo-bulks from the same scRNA-seq dataset that was also used as reference for the method. As we used data from two different organisms, we adapted the ‘species’ parameter accordingly. Other parameters remained with their default values as implemented in omnideconv. As recommended by the authors, we used un-normalized count values for scRNA-seq and bulk RNA-seq samples.

### Bisque

The R package BisqueRNA v1.0.5 was forked from https://github.com/cozygene/bisque and installed via the omnideconv package. We kept all parameters at their default as implemented in omnideconv and provided un-normalized count values for scRNA-seq and bulk RNA-seq samples. Bisque will then internally normalize both input data matrices to CPM counts by default.

### CIBERSORTx

We used the ‘fractions’ Docker image of CIBERSORTx (version 1.0 from 12/21/2019), which can be downloaded from the https://cibersortx.stanford.edu/ website. CIBERSORTx has two options for batch correction, S-mode and B-mode; we did not apply any batch correction in our analyses, as both correction modes yielded suboptimal results, particularly S-mode. Other parameters remained with their default values as implemented in omnideconv. As recommended by the authors, we used TPM normalized values for bulk RNA-seq samples and CPM normalization for the scRNA-seq samples.

### DWLS

We originally cloned the R package DWLS v0.1 from https://bitbucket.org/yuanlab/dwls/src/master/ and installed it via omnideconv. The original DWLS exhibited prolonged runtimes during the construction of the signature matrix. A primary reason for this was the absence of parallel processing in the original DWLS, rendering it incompatible with multicore machines. To address this, we introduced a new parameter, ’ncores,’ which allows users to specify the number of cores to be utilized. This parameter activates parallelization for MAST[84] functions like MAST::zlm, and enables us to leverage parallelized versions of iterative functions (e.g., replacing ’lapply’ with ’mclapply’) for enhanced parallel utilization.

In addition to parallelization, we conducted profiling to pinpoint performance bottlenecks within the DWLS program. Our analysis identified three key functions responsible for extended runtimes:

1. The internal method ’stat.log2.’
2. MAST::fromFlatDF, which was subsequently replaced with MAST::fromMatrix for streamlined data handling and improved speed.
3. The internal method ’m.auc,’ which, upon evaluation, was found to be unused for signature creation and was subsequently removed.

These optimizations not only significantly improved runtimes but also simplified the program structure. For example, reducing unnecessary dataframe creations in the ’stat.log2’ function minimized memory assignments, resulting in reduced code complexity and faster execution. The overall outcome of this improved DWLS implementation is a 3x speedup on a multicore machine with 10 cores, and a 2x improvement running DWLS with a single core.

The optimized version of DWLS can be executed in *omnideconv* with the parameter

*dwls_method* set to *mast_optimized*, which is the default for all results presented in this work.

Finally, we used DWLS with default parameters, only changing the pval_cutoff parameter to 0.05.

### MuSiC

The R package MuSiC v0.3.0 was forked from https://github.com/xuranw/MuSiC and installed via omnideconv. While a version 1.0.0 is already available, the only difference between the versions is the additional support for SingleCellExperiment data objects. The method includes information of patients in its deconvolution algorithm, which we added via the ‘batch_ids’ parameter of the ‘deconvolute()’ function in omnideconv. All parameters were set to default as implemented in omnideconv, scRNA-seq expression values were left un-normalized and bulk RNA-seq values were TPM normalized.

### Scaden

The python package Scaden v1.1.2 was forked from https://github.com/KevinMenden/scaden. All parameters were set to default as implemented in omnideconv, scRNA-seq expression values were left un-normalized and bulk RNA-seq values were TPM normalized. Scaden first trains its internal deep learning model by simulating pseudo-bulk data from the provided reference dataset and then uses this model to predict actual cell-type fractions. We left the parameters for the model building at the default values (5000 training steps, batch size of 128, learning rate of 1e-4 and 1000 pseudo-bulk samples of 100 cells each).

### SCDC

The R package SCDC v0.0.0.9 was forked from https://github.com/meichendong/SCDC and installed via omnideconv. Parameters for deconvolution were set to default as implemented in omnideconv, scRNA-seq expression values were left un-normalized and bulk RNA-seq values were TPM normalized. In contrast with other methods, SCDC allows the usage of multiple single cell RNAseq references. In this study, the method was used with just one single cell dataset as reference to keep the benchmarking as close as possible across methods.

## Declarations

### Ethics approval and consent to participate

Not applicable.

### Consent for publication

Not applicable.

### Availability of data and materials

All bulk datasets that are part of this study can be accessed via *deconvData* at https://figshare.com/projects/deconvData/197794; it includes raw and TPM-normalized gene expression matrices and ground truth annotations for each real and simulated sample. In addition, we give access to the processed counts and annotations of every scRNA-seq dataset that was used in this study.

A Docker image that includes an installed version of omnideconv and its dependencies can be accessed via Dockerhub at https://hub.docker.com/repository/docker/alexd13/omnideconv_benchmark/general, and a compressed file of the same image (version 2.0) is located in the *deconvData* repository to provide also long-term support.

The omnideconv ecosystem and all its included software packages are available at https://omnideconv.org/. It includes the *omnideconv* R package (https://github.com/omnideconv/omnideconv), the *deconvExplorer* web server (https://daisybio.ls.tum.de/deconvexplorer/) and the *deconvData* data access (https://figshare.com/projects/deconvData/197794). The code to run *deconvBench*, reproduce the figures and simulation scenarios and a detailed documentation of *deconvBench* can be found at https://github.com/omnideconv/deconvBench.

### Competing interests

FF consults for iOnctura. GS is an employee of Boehringer Ingelheim International Pharma GmbH & Co KG, Biberach, Germany. ML consults for mbiomics GmbH.

## Funding

This work was supported by European Cooperation in Science and Technology (COST) Action “Mye-InfoBank” (CA20117, supported by the EU Framework Program Horizon 2020), by the de.NBI Cloud within the German Network for Bioinformatics Infrastructure (de.NBI), and by ELIXIR-DE (Forschungszentrum Jülich and W-de.NBI-001, W-de.NBI-004, W-de.NBI-008, W-de.NBI-010, W-de.NBI-013, W-de.NBI-014, W-de.NBI-016, W-de.NBI-022). FF was supported by the Austrian Science Fund (FWF) (no. T 974-B30 and FG 2500-B) and by the Oesterreichische Nationalbank (OeNB) (no. 18496). FE was supported by the Austrian Science Fund (FWF) (Special Research Program F7804-B and I5184). ML and AD were supported by the German Federal Ministry of Education and Research (BMBF) within the framework of the CompLS research and funding concept [031L0294B (NetfLID)]. The computational results presented here have been achieved in part using the LEO HPC infrastructure of the University of Innsbruck and HPC infrastructure funded by the Deutsche Forschungsgemeinschaft (DFG, German Research Foundation) [422216132].

## Authors’ contributions

**Conceptualization:** ML, FF

**Data curation:** AD, LM, GS, FF1

**Formal analysis:** AD, LM

**Funding acquisition:** FE, ML, FF

**Investigation:** AD, LM, FF, ML

**Methodology:** AD, LM, KP, BE, CZ, KR, GS, FM

Project administration: **ML, FF**

**Resources:** FE, ML, FF

**Software:** AD, LM, GS, KP, BE, CZ, KR, FM

**Supervision:** FE, ML, FF, FM

**Validation:** AD, LM, KP, BE

**Visualization:** AD, LM

**Writing – original draft:** AD, LM, ML, FF

**Writing – review & editing:** AD, LM, BE, GS, FM, ML, FF

## Acknowledgements

We thank Nicolas Goedert and Yuyu Liang for their active development of a first version of the omnideconv package.

## Notes

https://figshare.com/projects/deconvData/197794

### Summary of Updates

One method (CIBERSORTx) showed different performances when using its 'S-mode' batch correction with the webtool version versus the Docker version. We have now disabled this batch correction entirely throughout all analyses.

