## Supplementary for "omnideconv: a unifying framework for using and benchmarking single-cell-informed deconvolution of bulk RNA-seq data"

### Supplementary Figures

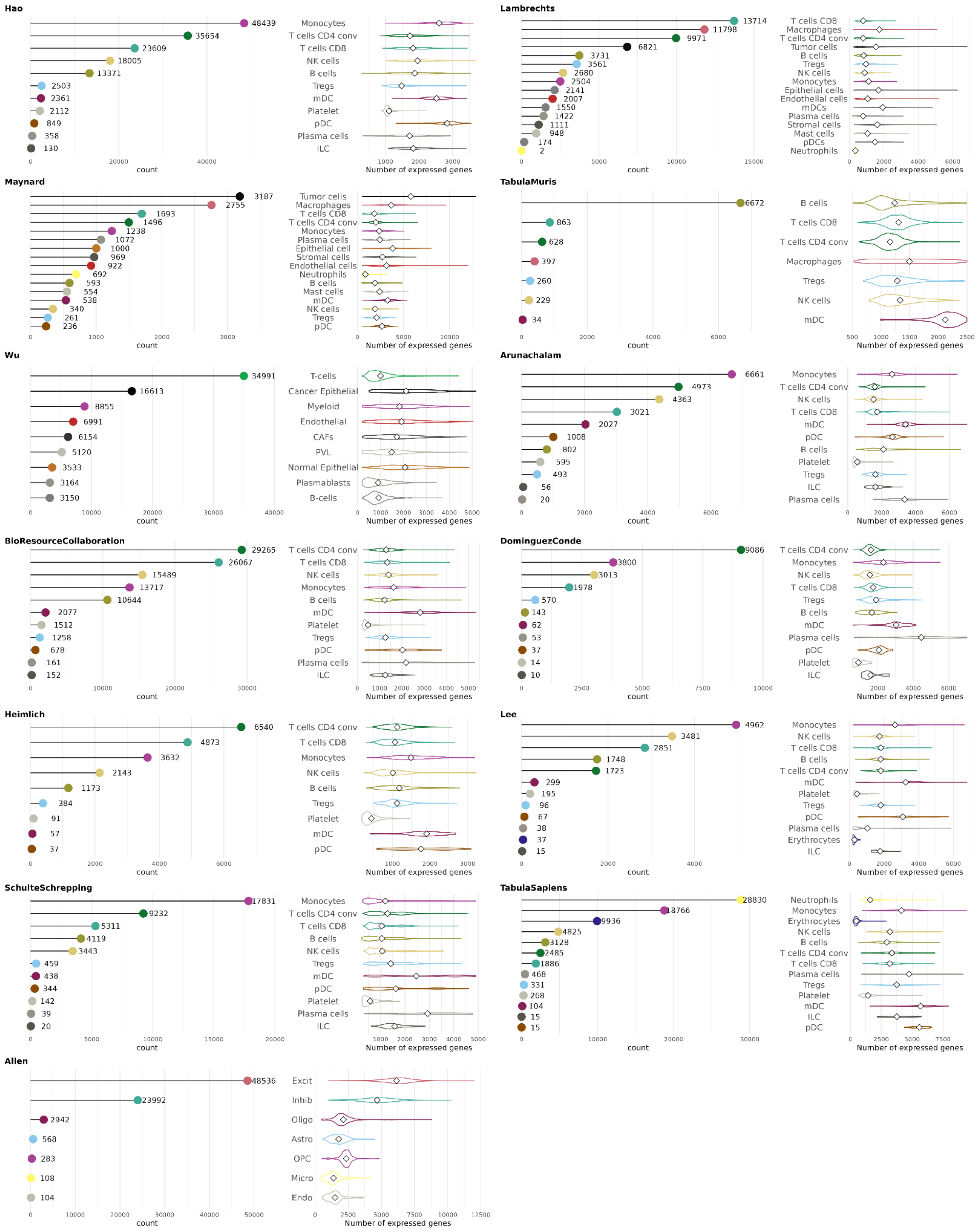

**Figure S1:** Overview of scRNA-seq datasets used in this benchmark. Displayed are the distribution of expression values in each cell type (right panel, respectively) and the number of cells for each cell type (left panel, respectively).

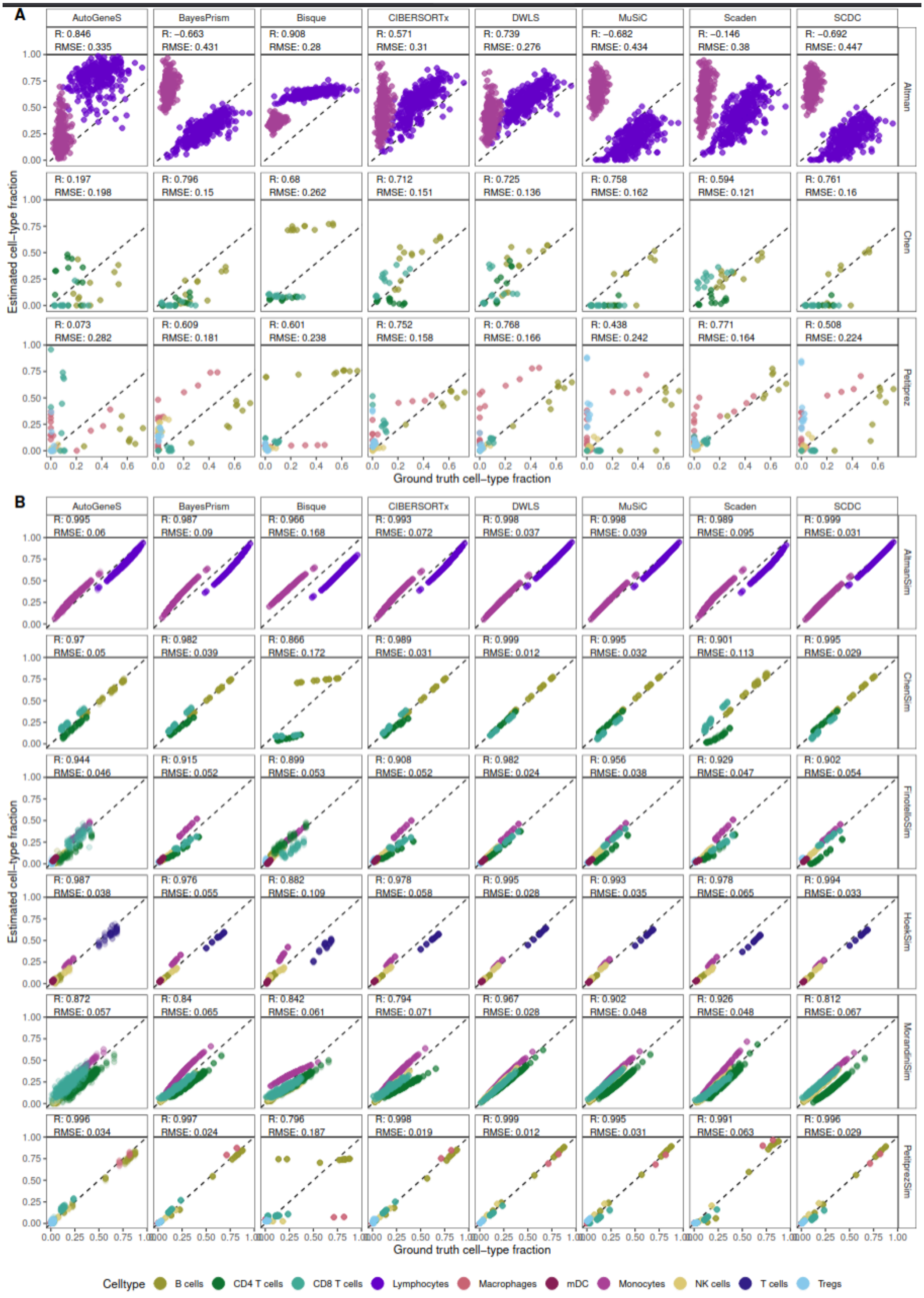
A
AutoGeneS
BayesPrism
Bisque
CIBERSORTx
DWLS
MuSiC
Scaden
SCDC
Altman
R: 0.846 RMSE: 0.335
R: −0.663 RMSE: 0.431
R: 0.908 RMSE: 0.28
R: 0.571 RMSE: 0.31
R: 0.739 RMSE: 0.276
R: −0.682 RMSE: 0.434
R: −0.146 RMSE: 0.38
R: −0.692 RMSE: 0.447
Chen
R: 0.197 RMSE: 0.198
R: 0.796 RMSE: 0.15
R: 0.68 RMSE: 0.262
R: 0.712 RMSE: 0.151
R: 0.725 RMSE: 0.136
R: 0.758 RMSE: 0.162
R: 0.594 RMSE: 0.121
R: 0.761 RMSE: 0.16
Petitprez
R: 0.073 RMSE: 0.282
R: 0.609 RMSE: 0.181
R: 0.601 RMSE: 0.238
R: 0.752 RMSE: 0.158
R: 0.768 RMSE: 0.166
R: 0.438 RMSE: 0.242
R: 0.771 RMSE: 0.164
R: 0.508 RMSE: 0.224
Estimated cell−type fraction
Ground truth cell−type fraction
B
AltmanSim
R: 0.995 RMSE: 0.06
R: 0.987 RMSE: 0.09
R: 0.966 RMSE: 0.168
R: 0.993 RMSE: 0.072
R: 0.998 RMSE: 0.037
R: 0.998 RMSE: 0.039
R: 0.989 RMSE: 0.095
R: 0.999 RMSE: 0.031
ChenSim
R: 0.97 RMSE: 0.05
R: 0.982 RMSE: 0.039
R: 0.866 RMSE: 0.172
R: 0.989 RMSE: 0.031
R: 0.999 RMSE: 0.012
R: 0.995 RMSE: 0.032
R: 0.901 RMSE: 0.113
R: 0.995 RMSE: 0.029
FinotelloSim
R: 0.944 RMSE: 0.046
R: 0.915 RMSE: 0.052
R: 0.899 RMSE: 0.053
R: 0.908 RMSE: 0.052
R: 0.982 RMSE: 0.024
R: 0.956 RMSE: 0.038
R: 0.929 RMSE: 0.047
R: 0.902 RMSE: 0.054
HoekSim
R: 0.987 RMSE: 0.038
R: 0.976 RMSE: 0.055
R: 0.882 RMSE: 0.109
R: 0.978 RMSE: 0.058
R: 0.995 RMSE: 0.028
R: 0.993 RMSE: 0.035
R: 0.978 RMSE: 0.065
R: 0.994 RMSE: 0.033
MorandiniSim
R: 0.872 RMSE: 0.057
R: 0.84 RMSE: 0.065
R: 0.842 RMSE: 0.061
R: 0.794 RMSE: 0.071
R: 0.967 RMSE: 0.028
R: 0.902 RMSE: 0.048
R: 0.926 RMSE: 0.048
R: 0.812 RMSE: 0.067
PetitprezSim
R: 0.996 RMSE: 0.034
R: 0.997 RMSE: 0.024
R: 0.796 RMSE: 0.187
R: 0.998 RMSE: 0.019
R: 0.999 RMSE: 0.012
R: 0.995 RMSE: 0.031
R: 0.991 RMSE: 0.063
R: 0.996 RMSE: 0.029
Estimated cell−type fraction
Ground truth cell−type fraction
Celltype
B cells
CD4 T cells
CD8 T cells
Lymphocytes
Macrophages
mDC
Monocytes
NK cells
T cells
Tregs

**Figure S2:** As Figure 2, but for (**A**) real bulk datasets *Altman*[6], *Petitprez*[7] and *Chen*[8] and (**B**) the pseudobulk representations *AltmanSim*, *ChenSim, FinotelloSim, HoekSim, MorandiniSim, PetitprezSim* (see Methods for details on simulation). The *HaoSub* dataset was used as a reference for human datasets, just as in Figure 2. We show results for mouse bulk and pseudo-bulk data in panels A and B, with a reference from the Tabula Muris (*TM)* dataset.

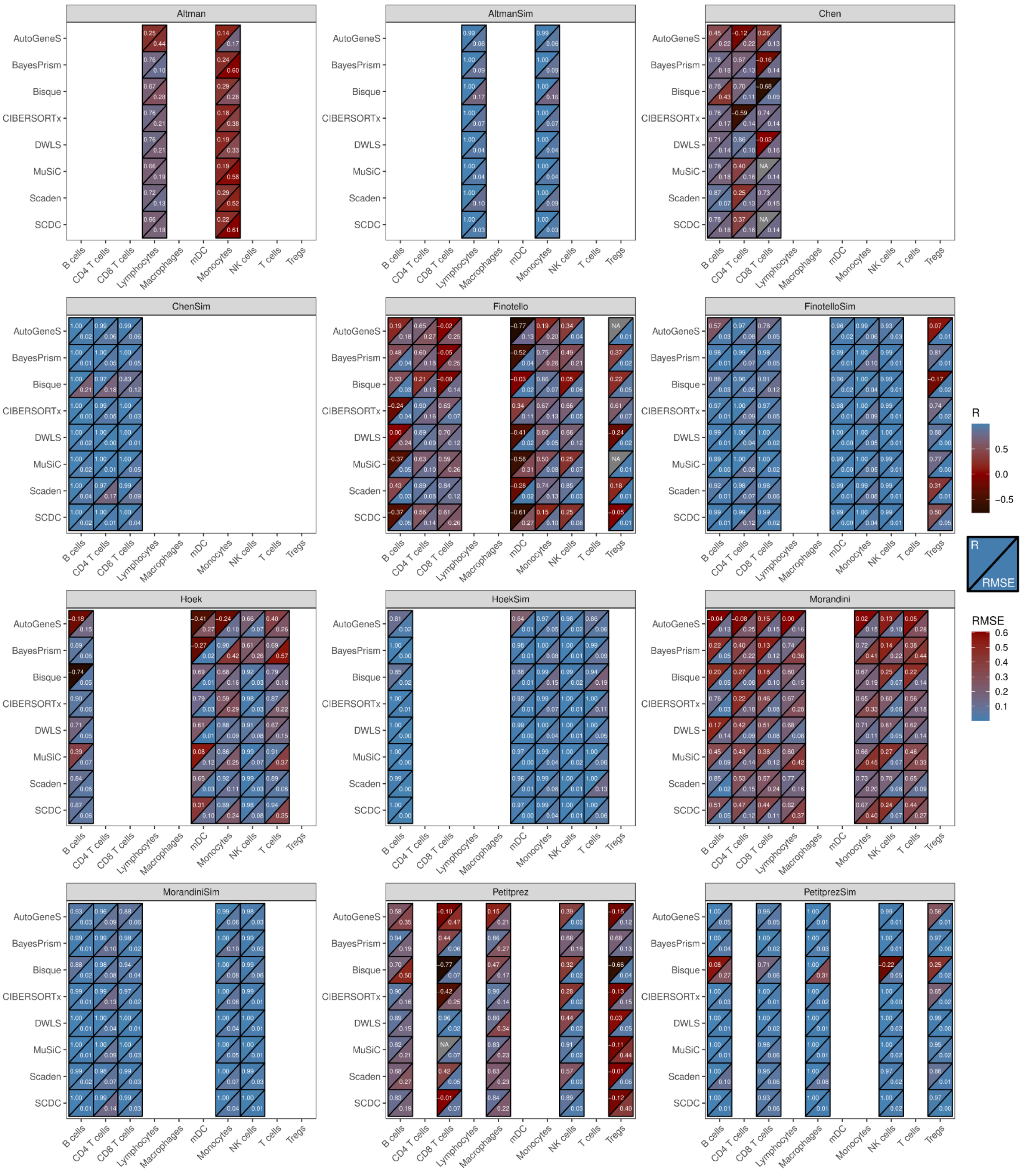

**Figure S3:** Pearson correlation coefficient (upper triangles) and RMSE (lower triangles) for eight deconvolution methods in twelve datasets (real bulk and pseudobulk, six each) of two different organisms: human (*Altman, AltmanSim, Finotello, FinotelloSim, Hoek, HoekSim, Morandini, MorandiniSim*) and mouse (*Chen, ChenSim, Petitprez, PetitprezSim*). The *HaoSub* dataset was used as a reference for human datasets, the Tabula Muris (*TM)* dataset for mouse datasets.

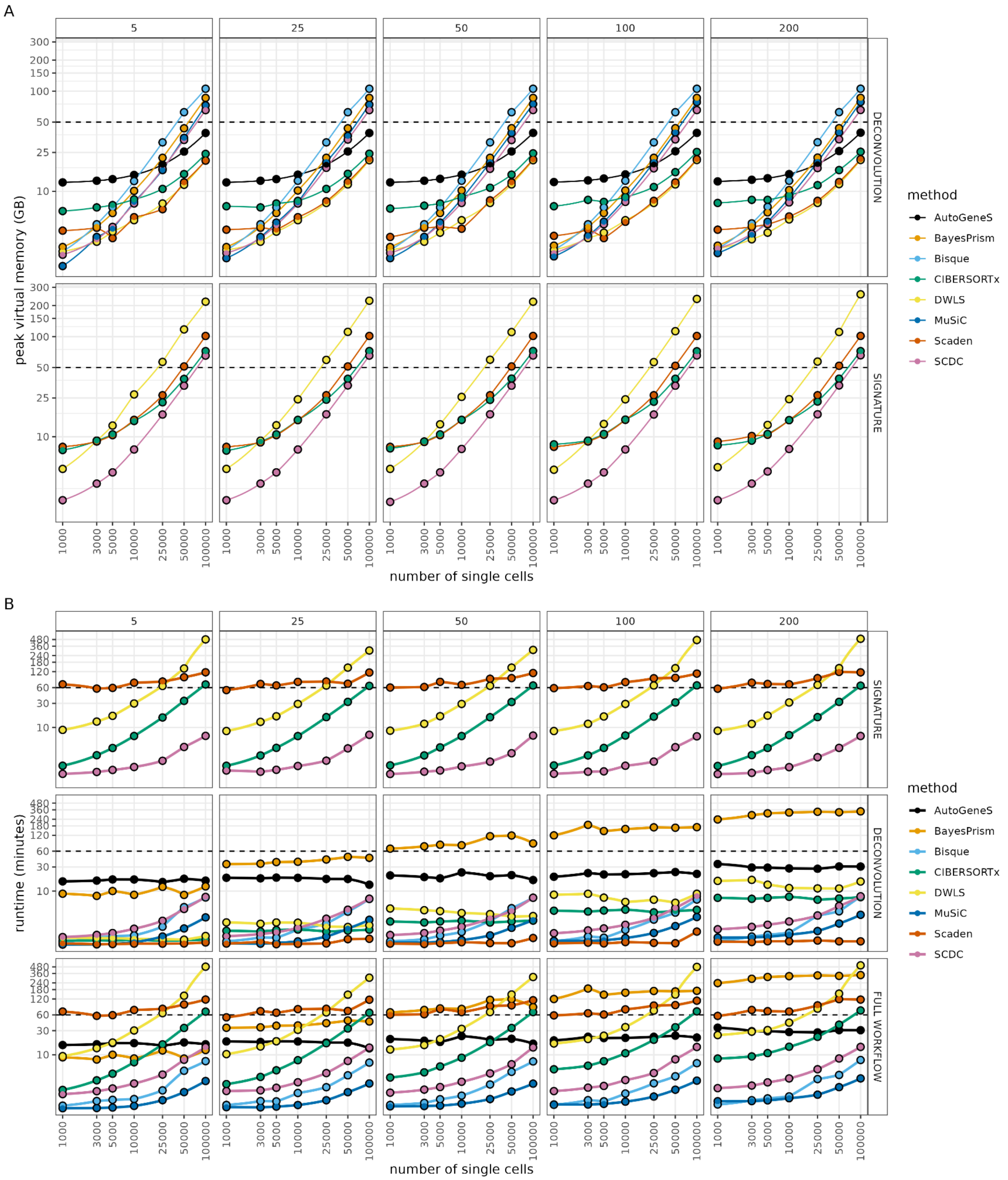

**Figure S4:** Resources required by deconvolution methods in terms of peak virtual memory (**A**) and runtime (**B**). Each method was supplied with increasingly larger single-cell reference datasets (x-axis) and was tasked to deconvolve an increasing number of pseudobulk samples (columns). For methods that calculate their signature in a separate step, the resource usage was measured separately for signature creation and follow-up deconvolution. In case of the runtime, the values for these two steps were also summed up to display the runtime of a complete workflow (rows). Note the log scaling of both the x and y axis in order to avoid overplotting.

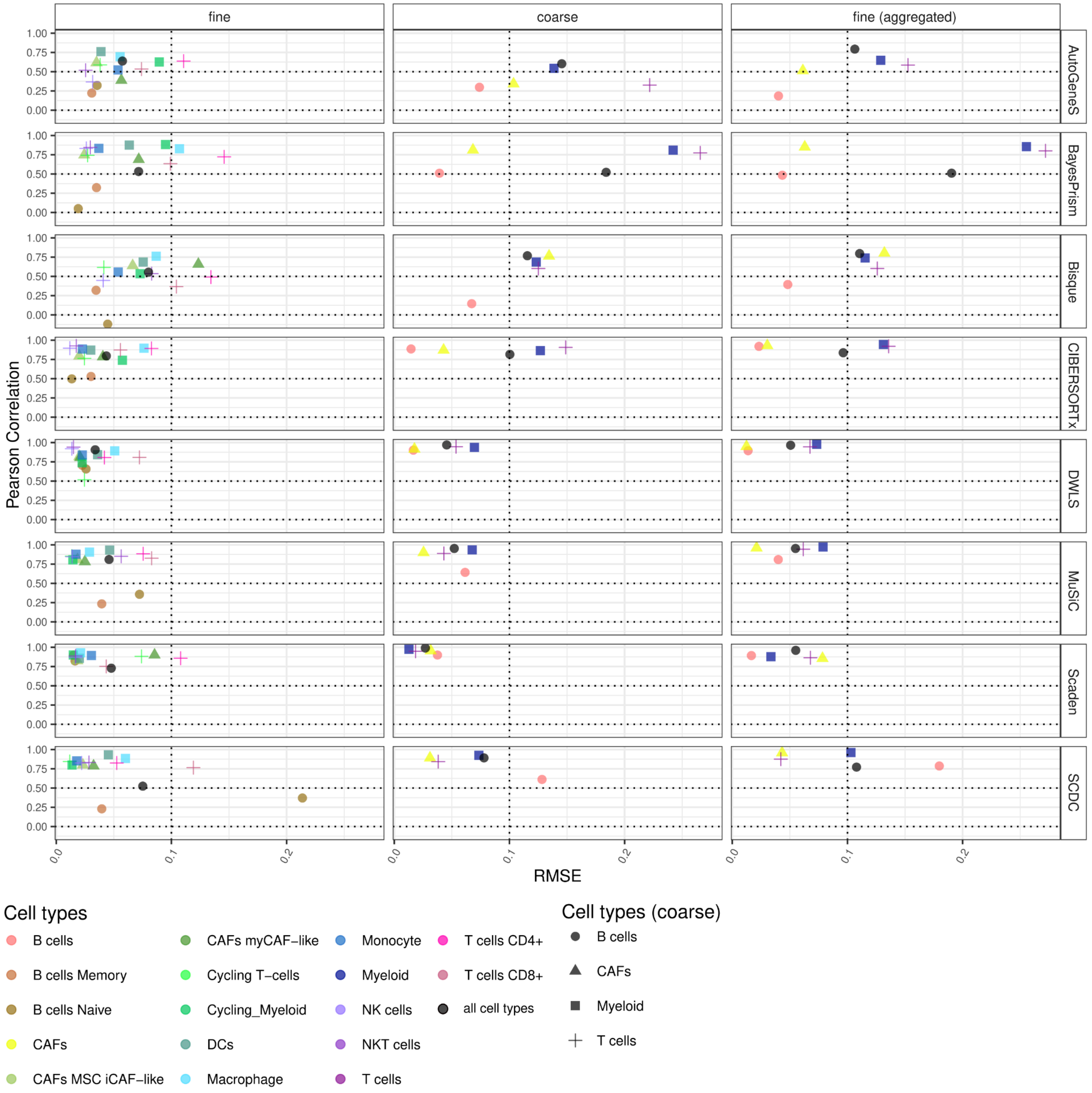

**Figure S5: Method performances with different annotation granularities of the reference, *Wu* dataset.**
**(A)** Pearson correlation coefficient and RMSE values computed for the cell type-specific estimates obtained on pseudo-bulks (n=50) simulated from the *Wu* dataset for fine and coarse annotation levels, and fine aggregated annotation level. Celltype abbreviations: cancer-associated fibroblasts (CAFs), mesenchymal-derived cancer-associated inflammatory-like fibroblasts (CAFs MSC iCAF-like), cancer-associated fibroblasts myofibroblastic-like (CAFS myCAF-like), dendritic cells (DCs), natural killer cells (NK cells), T cells NK-like (NK T cells).

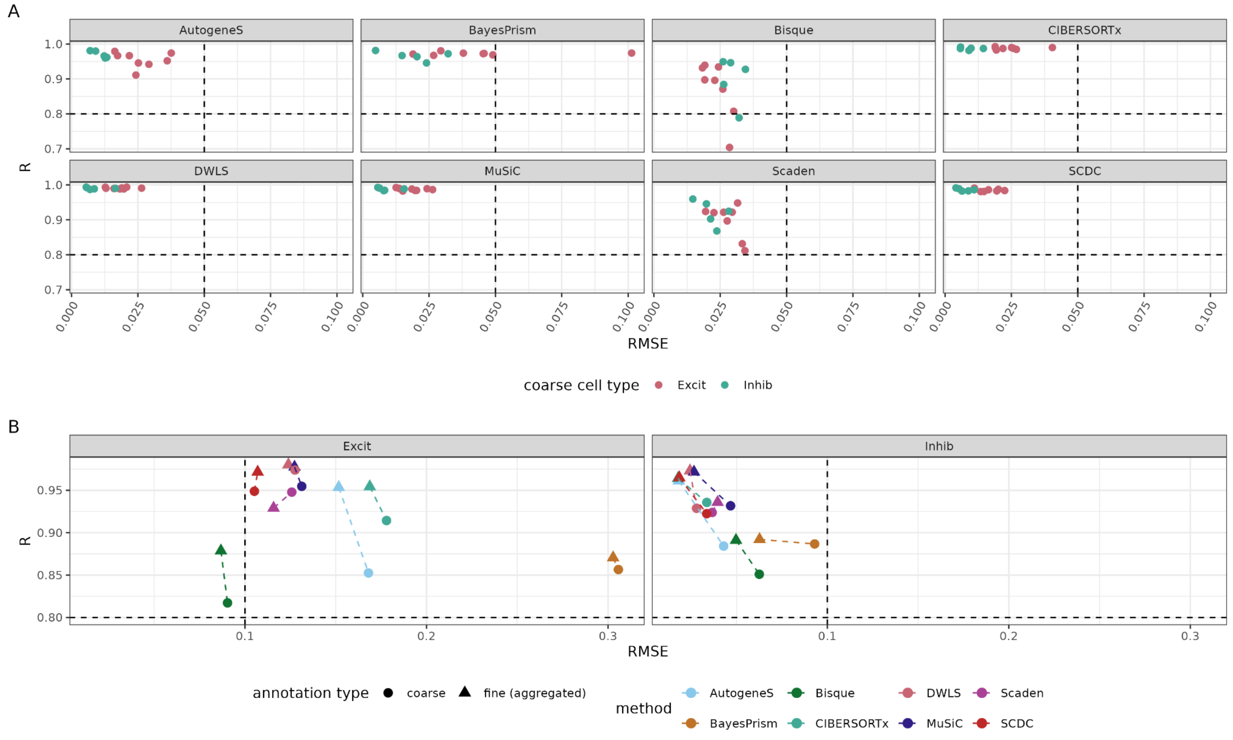

**Figure S6: Method performances with different annotation granularities of the reference, *Allen* dataset**
**(A)** Pearson correlation coefficient and RMSE values computed for the estimates of subtypes of Inhibitory (Inhib) and Excitatory (Excit) neurons, obtained on pseudo-bulks (n=50) simulated from the *Allen* dataset for fine annotation level. The points are colored according to the coarse annotation. (B) Pearson correlation coefficient and RMSE values computed for the estimates of Inhibitory (Inhib) and Excitatory (Excit) neurons, obtained on pseudo-bulks (n=50) simulated from the *Allen* dataset for coarse and fine aggregated annotation levels. The points are colored according to the method.

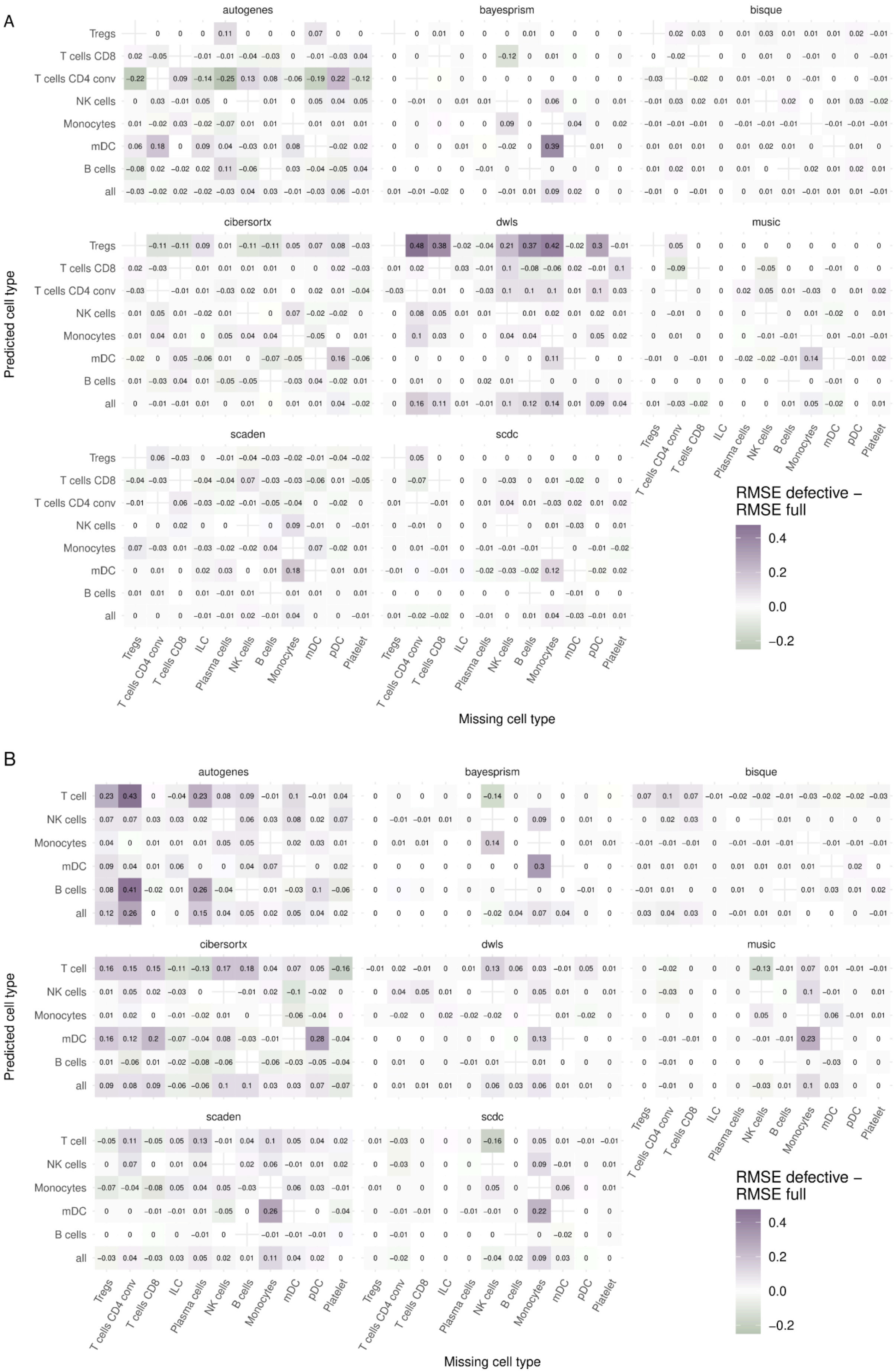
A
autogenes
bayesprism
bisque
cibersortx
dwls
music
scaden
scdc
Tregs
T cells CD8
T cells CD4 conv
NK cells
Monocytes
mDC
B cells
all
Predicted cell type
Missing cell type
ILC
Plasma cells
pDC
Platelet
RMSE defective – RMSE full
0.4
0.2
0.0
−0.2
B
T cell

**Figure S7:** Difference between the Root mean squared error (RMSE) between cell type predictions obtained with methods trained on the defective *HaoSub* reference (i.e. with one missing cell type, indicated in columns) and cell type ground truth fractions, and RMSE obtained with the methods trained on the full reference and cell type ground truth fractions for the (**A**) *Finotello* (n=9) and (**B**) *Hoek* bulk dataset (n=8). A positive value means that the removal of a specific cell type worsens the deconvolution performance, and, vice versa, a negative value indicates improved performance.

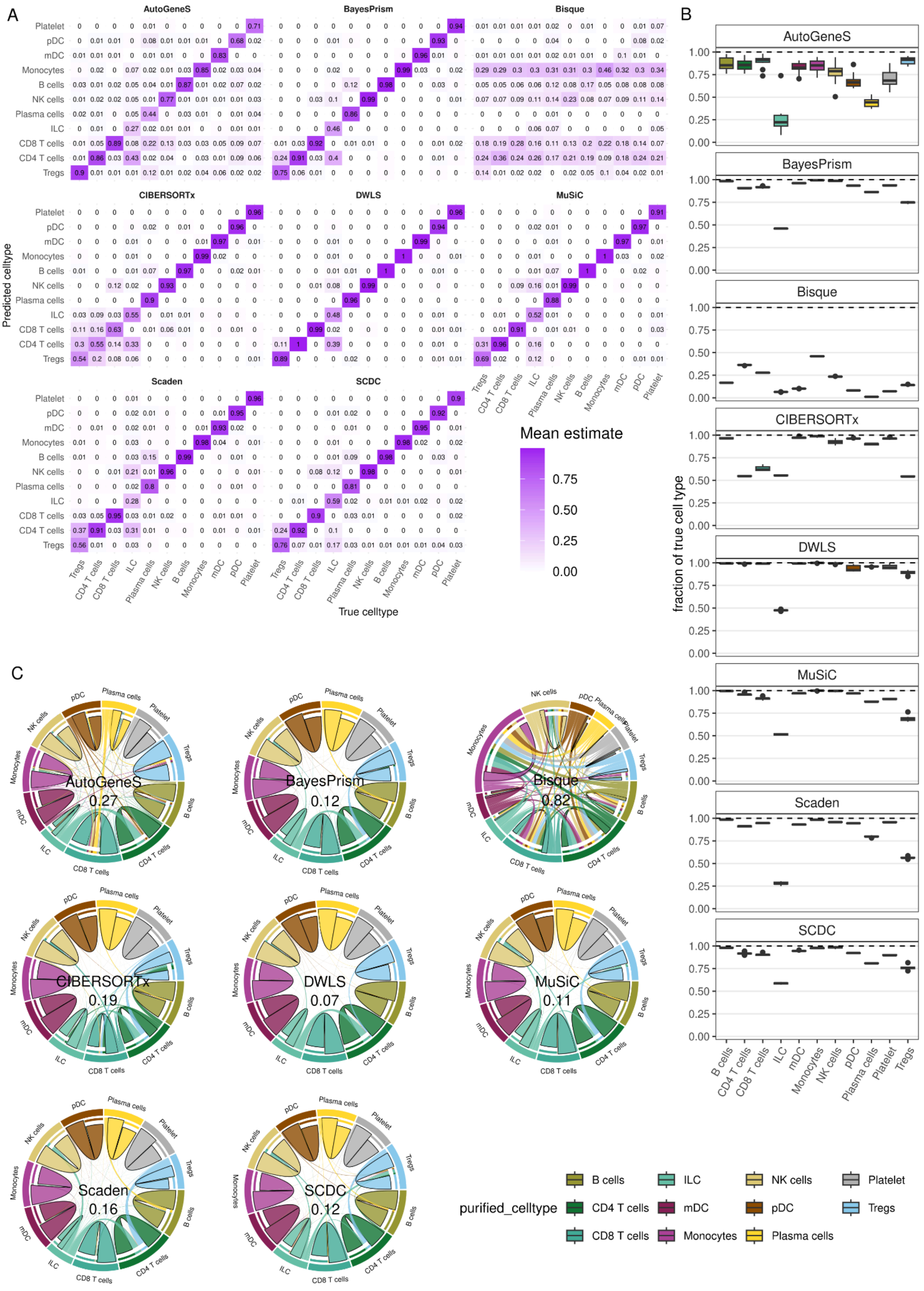
A
AutoGeneS
BayesPrism
Bisque
CIBERSORTx
DWLS
MuSiC
Scaden
SCDC
Predicted celltype
True celltype
Platelet
pDC
mDC
Monocytes
B cells
NK cells
Plasma cells
ILC
CD8 T cells
CD4 T cells
Tregs
Mean estimate
0.75
0.50
0.25
0.00
B
fraction of true cell type
1.00
0.75
0.50
0.25
0.00
C
AutoGeneS 0.27
BayesPrism 0.12
Bisque 0.82
CIBERSORTx 0.19
DWLS 0.07
MuSiC 0.11
Scaden 0.16
SCDC 0.12
purified_celltype
B cells
CD4 T cells
CD8 T cells
ILC
mDC
Monocytes
NK cells
pDC
Plasma cells
Platelet
Tregs

**Figure S8:** Results of the spillover analysis with pseudo-bulks generated from the *Hao* scRNA-seq dataset with the *HaoSub* dataset used as deconvolution reference (see Methods for details). Each method was applied to simulated samples that contained only cells of one of eleven cell types. These samples (n=550) were then deconvolved and the average prediction for each cell type was considered. (**A**) displays the mean fraction predicted across samples, for each cell type. (**B**) Percentage of correctly predicted cell type abundance in the spillover analysis. The presented values correspond to the sum of the correctly predicted cell fractions for each method. **(C)** Chord diagrams showing the cell type predictions for the spillover analysis, similar to Fig 5B. The values in the center indicate the total fraction of predictions attributed to wrong cell types.

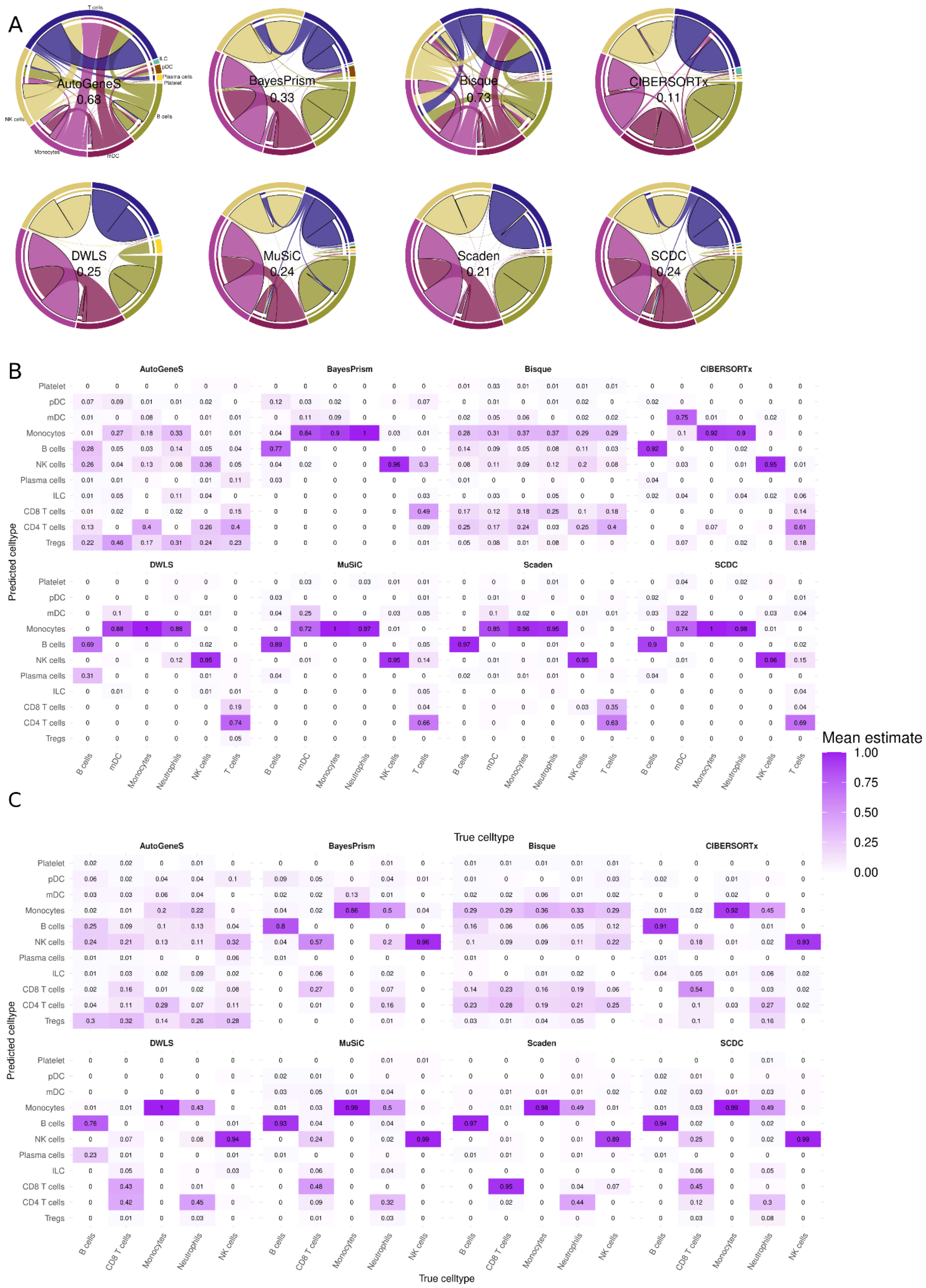
A
AutoGeneS
0.68
BayesPrism
0.33
Bisque
0.73
CIBERSORTx
0.11
DWLS
0.25
MuSiC
0.24
Scaden
0.21
SCDC
0.24
B
C
Predicted celltype
True celltype
Mean estimate
1.00
0.75
0.50
0.25
0.00
Platelet
pDC
mDC
Monocytes
B cells
NK cells
Plasma cells
ILC
CD8 T cells
CD4 T cells
Tregs
Neutrophils
T cells

**Figure S9: (A)** Chord diagrams showing the cell type predictions for the spillover analysis for the real purified *Hoek-pure* bulk dataset, similar to Figure 5B. The values in the center indicate the total fraction of predictions attributed to wrong cell types. The heatmaps below display the mean fraction predicted across samples, for each cell type, for the *Hoek-pure* **(B)** and *Linsley-pure* **(C)** real purified bulk datasets.
Note that for the deconvolution of both datasets, we used the *HaoSub* reference, which does not contain Neutrophils. However, there are purified Neutrophil samples in both *Hoek-pure* and *Linsley-pure* available.

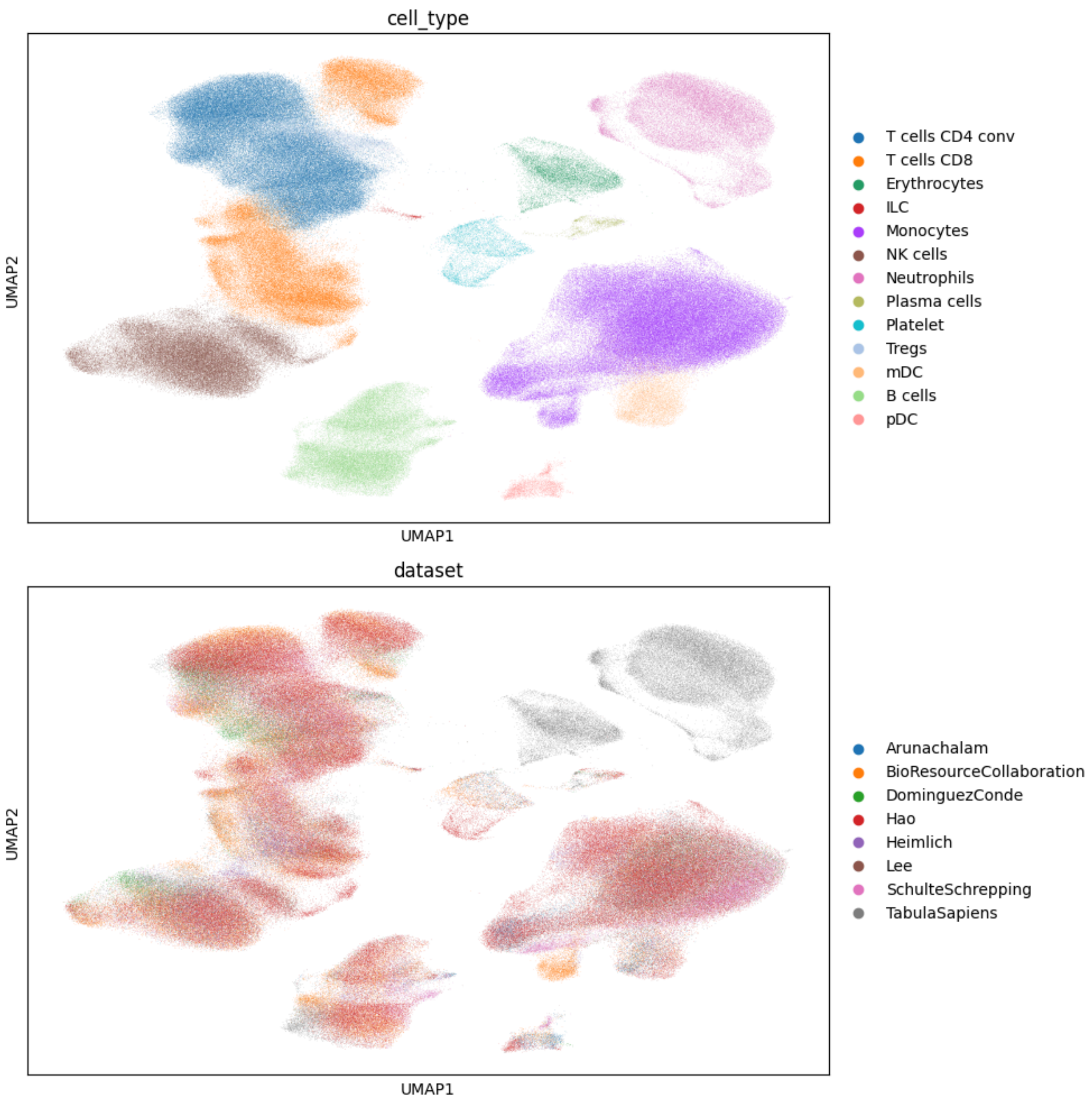

**Figure S10:** UMAP representation of the integration of eight scRNA-seq datasets (bottom) from healthy blood patients. Datasets were integrated using scANVI (Xu et al. 2021), where the batch variable was the individual donor IDs of the datasets, and our own manual coarse annotation was provided to scANVI for better integration performance. These coarse labels are based upon the original cell-type labels of the datasets. Finally, cells were labelled after successful integration using Leiden clustering and marker gene expression (top).

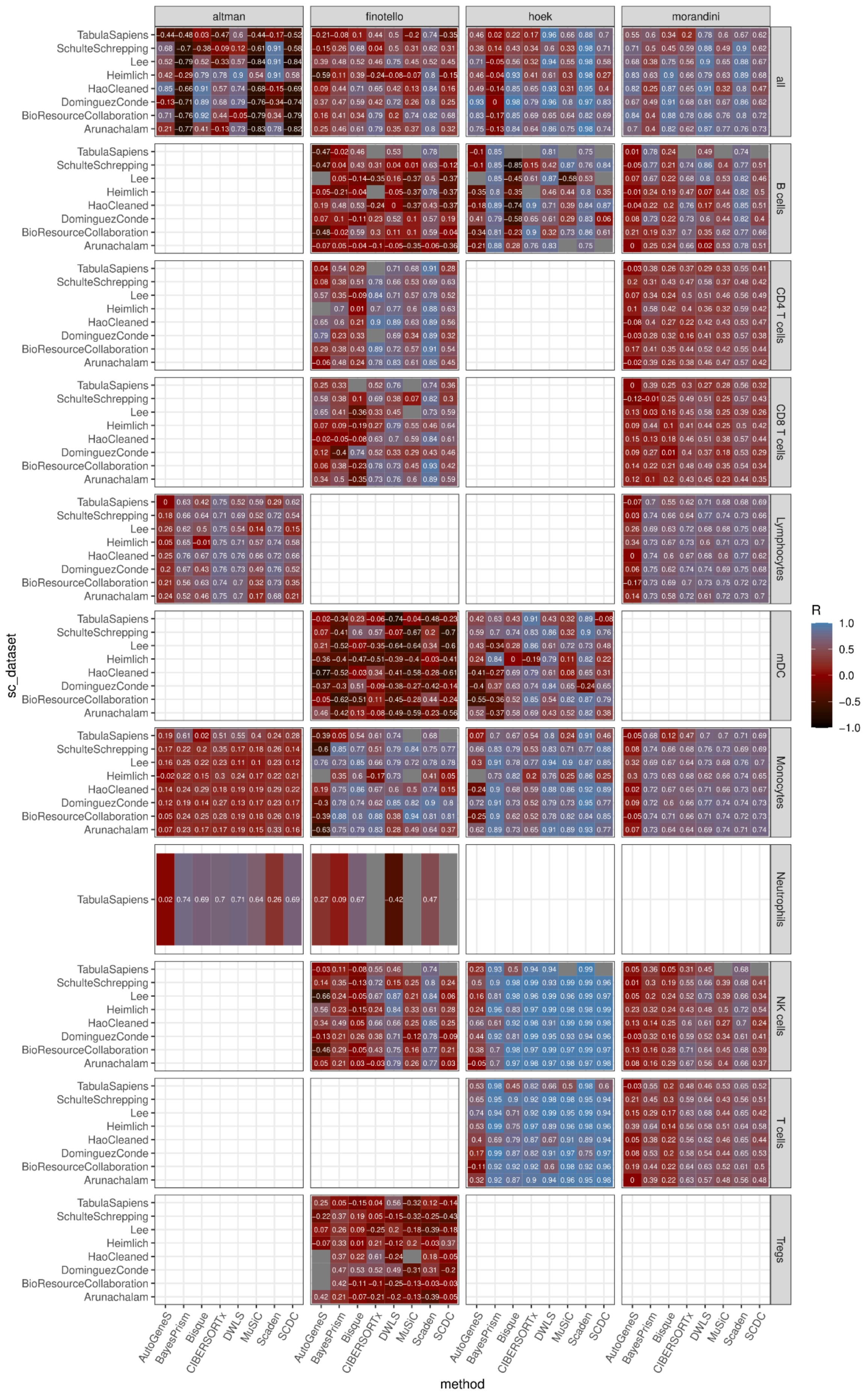

altman
finotello
hoek
morandini
all
B cells
CD4 T cells
CD8 T cells
Lymphocytes
mDC
Monocytes
Neutrophils
NK cells
T cells
Tregs
TabulaSapiens
SchulteSchrepping
Lee
Heimlich
HaoCleaned
DominguezConde
BioResourceCollaboration
Arunachalam
sc_dataset
AutoGeneS
BayesPrism
Bisque
CIBERSORTx
DWLS
MuSiC
Scaden
SCDC
method
R
1.0
0.5
0.0
−0.5
−1.0

**Figure S11:** Pearson correlation of cell-type predictions across eight different single-cell reference datasets (small rows) applied to four different bulk datasets (large columns). Deconvolution methods (small columns) provided estimates for different cell types (large rows), which were compared to FACS or CBC ground truth to calculate correlation. The large row with ‘all’ shows the correlation based on all cell-type estimates combined; Neutrophils could only be detected with one reference dataset (Tabula Sapiens).

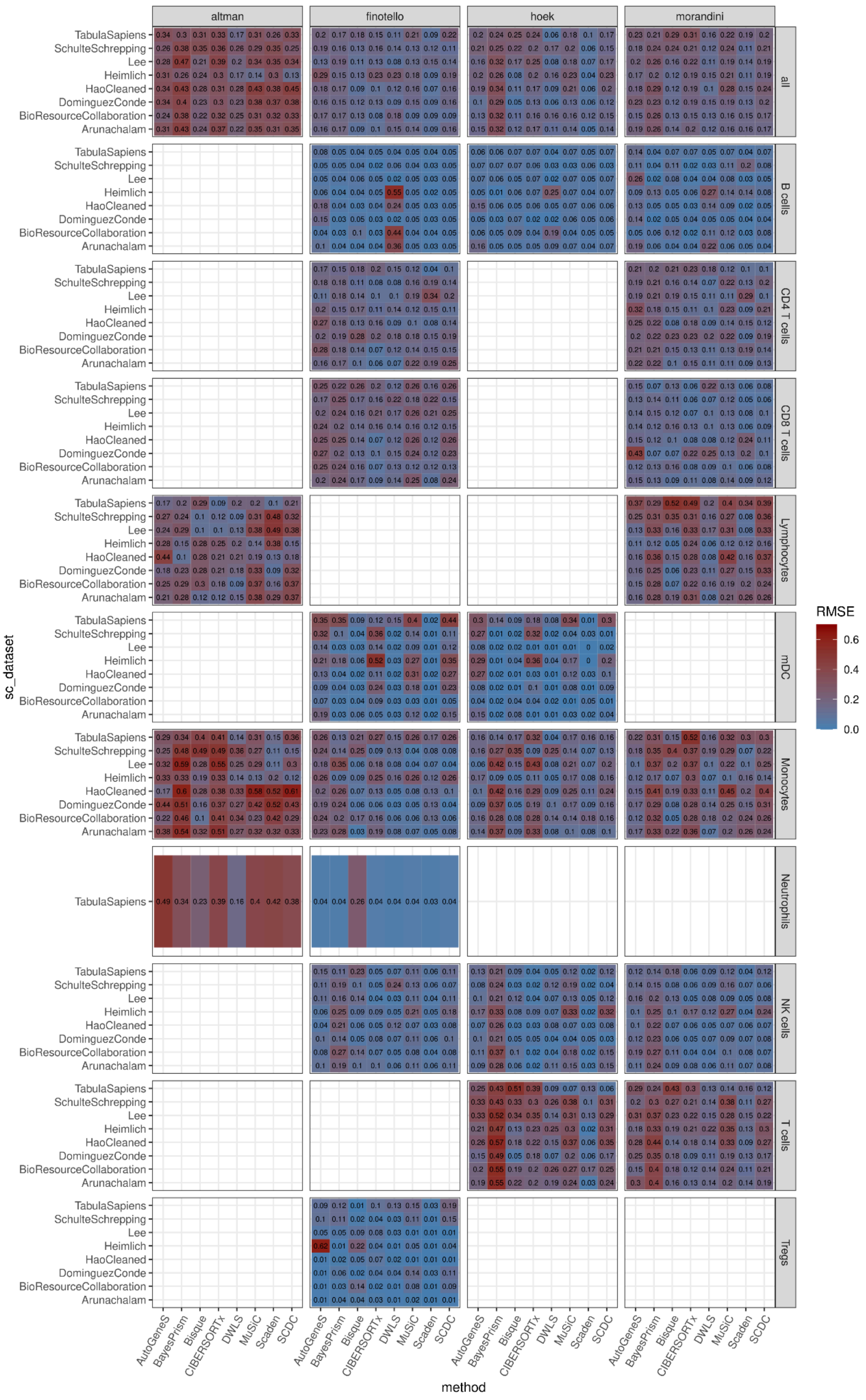

altman
finotello
hoek
morandini
all
B cells
CD4 T cells
CD8 T cells
Lymphocytes
mDC
Monocytes
Neutrophils
NK cells
T cells
Tregs
TabulaSapiens
SchulteSchrepping
Lee
Heimlich
HaoCleaned
DominguezConde
BioResourceCollaboration
Arunachalam
sc_dataset
AutoGeneS
BayesPrism
Bisque
CIBERSORTx
DWLS
MuSiC
Scaden
SCDC
method
RMSE
0.6
0.4
0.2
0.0

**Figure S12:** The same as Figure S12, but for root mean square error (RMSE).

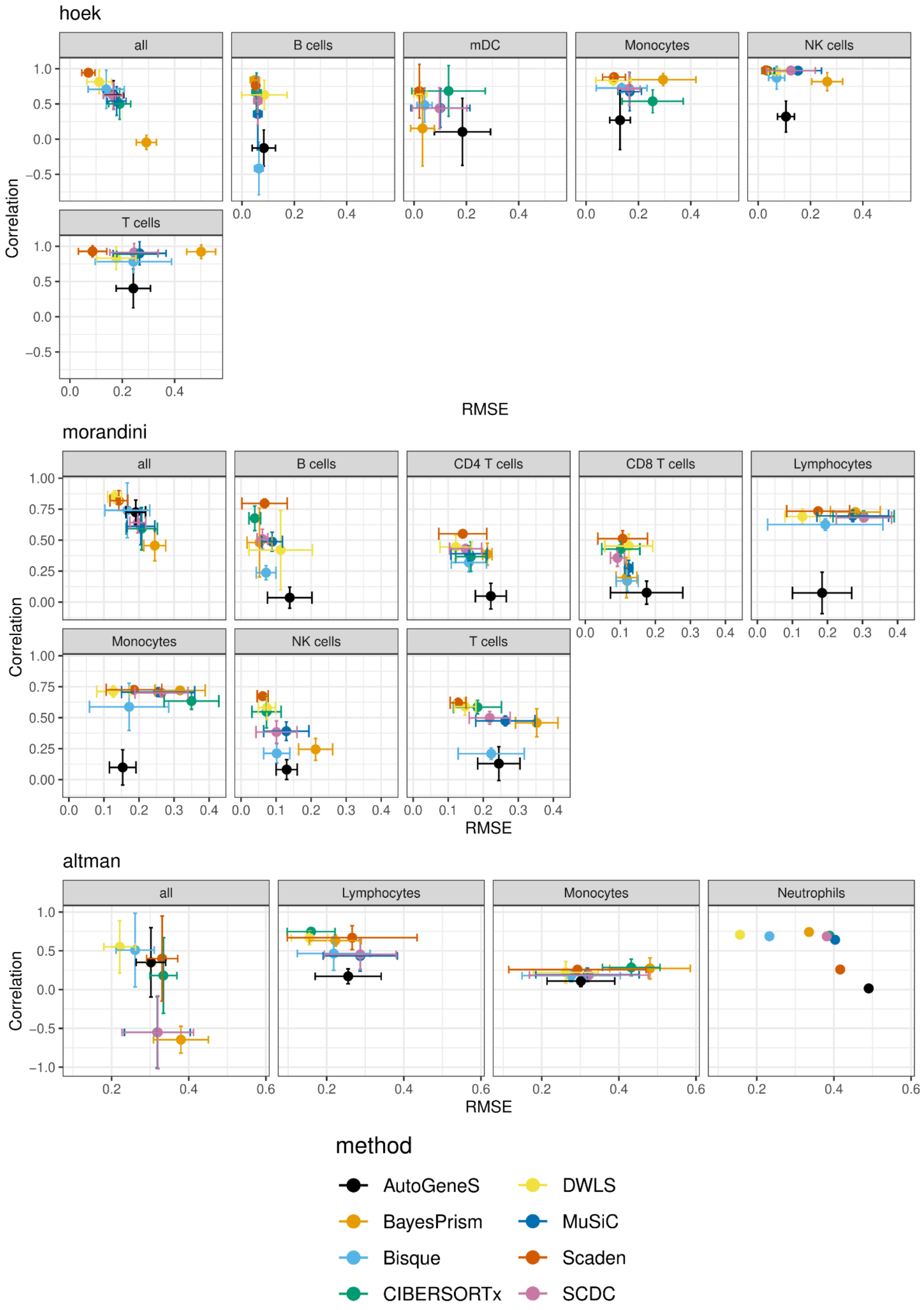
hoek
all
B cells
mDC
Monocytes
NK cells
T cells
Correlation
RMSE
morandini
all
B cells
CD4 T cells
CD8 T cells
Lymphocytes
Monocytes
NK cells
T cells
Correlation
RMSE
altman
all
Lymphocytes
Monocytes
Neutrophils
Correlation
RMSE
method
AutoGeneS
BayesPrism
Bisque
CIBERSORTx
DWLS
MuSiC
Scaden
SCDC

**Figure S13:** Mean Pearson correlation and mean RMSE for cell-type predictions across eight different single-cell reference datasets applied to the real *Hoek, Morandini* and *Altman* bulk datasets. Error bars represent the standard deviation of each method in a cell type, calculated across the eight different references. The box with ‘all’ shows the correlation and RMSE based on all cell-type fractions combined; Neutrophils could only be detected with one reference dataset (*Tabula Sapiens*).

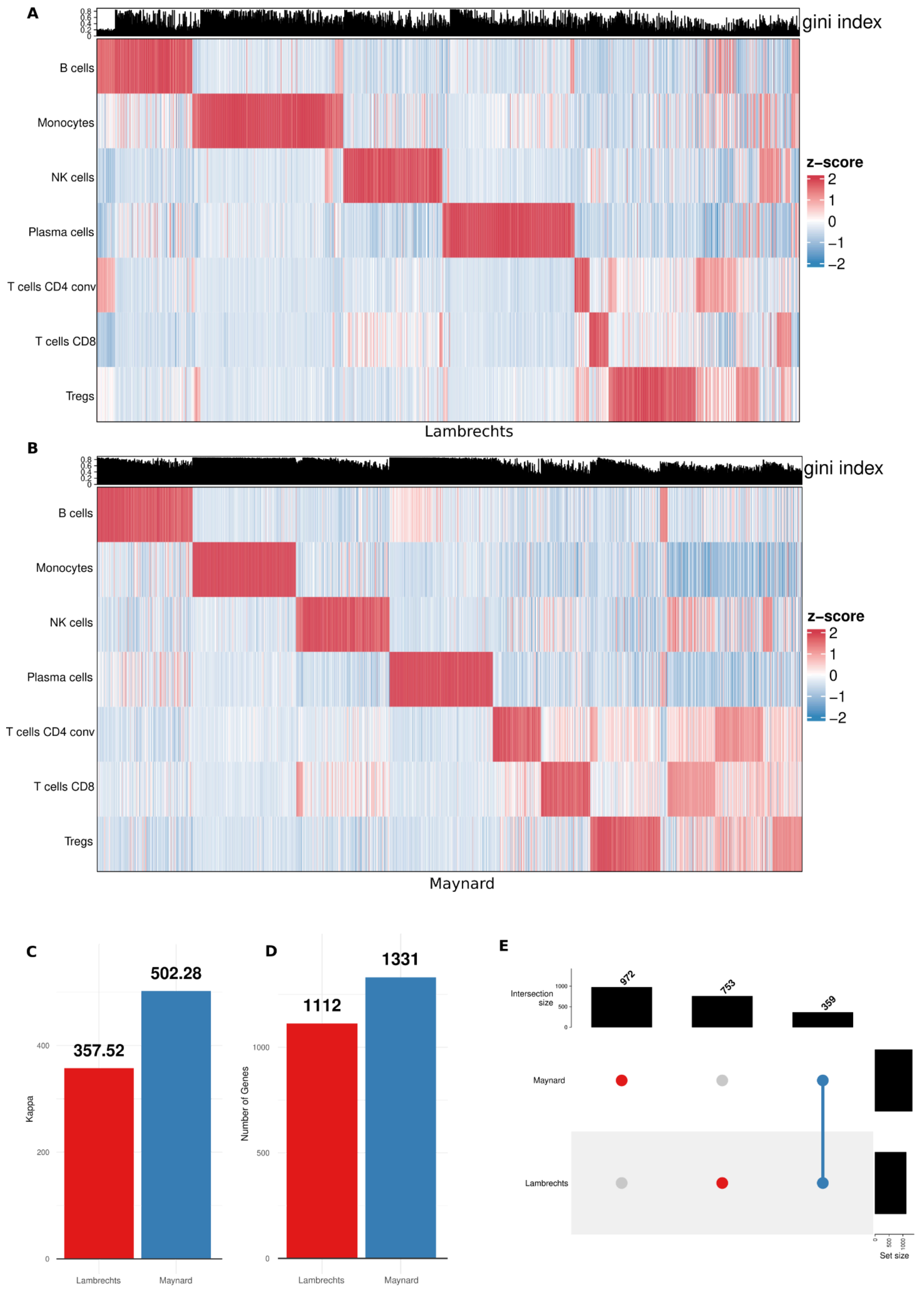
A
gini index
B cells
Monocytes
NK cells
Plasma cells
T cells CD4 conv
T cells CD8
Tregs
z–score
2
1
0
−1
−2
Lambrechts
B
gini index
B cells
Monocytes
NK cells
Plasma cells
T cells CD4 conv
T cells CD8
Tregs
z–score
2
1
0
−1
−2
Maynard
C
502.28
357.52
Kappa
400
200
0
Lambrechts
Maynard
D
1331
1112
Number of Genes
1000
500
0
Lambrechts
Maynard
E
Intersection size
972
753
359
Maynard
Lambrechts
Set size

**Figure S14:** *deconvExplorer* allows for manual signature exploration with a variety of visualizations; here, we show them for signatures created with DWLS and two different datasets (*Lambrechts*, *Maynard*), which have all been subset to contain seven intersecting cell types before being used as input for DWLS. (**A**) shows gene-wise z-scored heatmaps of the *Lambrechts* signature, and (**B**) shows the same for *Maynard*, where columns represent clustered genes. In addition, a barplot is shown on top of each heatmap, indicating the gene-wise Gini index. (**C-E**) display summary values for each signature: the condition number ("Kappa") (**C**) and number of genes that are part of the signature (**D**). Finally, the upset plot (**E**) shows the number of unique (red) and intersecting (blue) genes between the signatures, which can also be downloaded directly from *deconvExplorer*. See methods for a detailed description of the calculation of each value.

### Supplementary Tables

**Table S1:** List of all real and simulated bulk RNA-seq datasets used in this benchmarking study.

| name | tissue | organism | type of ground truth | n _samples | ground truth cell types | reference |
|---|---|---|---|---|---|---|
| Finotello | blood | human | FACS | 9 | NK cells, B cells, Tregs, mDC, Monocytes, Neutrophils, T cells CD8, T cells CD4 conv, Other | https://doi.org/10.1186/s13073-019-0638-6 |
| Hoek | blood | human | FACS | 8 | T cells, Monocytes, B cells, mDC, NK cells | https://doi.org/10.1371/journal.pone.0118528 |
| Hoek_purified | blood | human | purified populations | 48 | B cells, mDC, Monocytes, Neutrophils, NK cells, T cells | https://doi.org/10.1371/journal.pone.0118528 |
| Linsley_purified | blood | human | purified populations | 114 | B cells, Monocytes, Neutrophils, NK cells, T cells CD4, T cells CD8 | https://doi.org/10.1371/journal.pone.0109760 |
| Altman | blood | human | CBC | 322 | Basophils, Eosinophils, Lymphocytes, Monocytes, Neutrophils | https://doi.org/10.1038/s41590-019-0347-8 |
| Morandini | blood | human | FACS | 159 | Monocytes, Granulocytes, Lymphocytes, B cells, NK cells, T cells, T cells CD4, T cells CD8 | https://doi.org/10.1007/s11357-023-00986-0 |
| Vanderbilt | lung cancer | human | IHC | 8 | T cells CD4, T cells CD8 | https://doi.org/10.1186/s13073-019-0638-6 |
| Chen | spleen, lymph nodes, bone marrow, pbmc | mouse | FACS | 12 | Monocytes, B cells, T cells CD8, T cells CD4 conv | https://doi.org/10.1038/srep40508 |
| Petitprez | peritoneum, | mouse | FACS | 14 | T cells, T cells CD8, B cells, Tregs, NK cells, B cells germinal, Basophils, MAST cells, | https://doi.org/10.1186/s13073-020-00783-w |

| | | | | | |
|---|---|---|---|---|---|
| | spleen, blood, tumor | | | | Macrophages, Eosinophils, Neutrophils, Monocytes, B derived cells, Monocytes/Macrophages, NK/T cells, Granulocytes |
| Finotello-Sim | blood | human | pseudobulk (Hao, match FACS fractions) | 90 | Monocytes, T cells CD4 conv, T cells CD8, NK cells, B cells, Tregs, mDC |
| Hoek-Sim | blood | human | pseudobulk (Hao, match FACS fractions) | 80 | T cells, Monocytes, B cells, mDC, NK cells |
| Morandini-Sim | blood | human | pseudobulk (Hao, match FACS fractions) | 780 | Monocytes, B cells, NK cells, T cells CD4, T cells CD8 |
| Altman-Sim | blood | human | pseudobulk (Hao, match FACS fractions) | 1610 | Lymphocytes, Monocytes, Neutrophils |
| Chen-Sim | spleen, lymph nodes, bone marrow, pbmc | mouse | pseudobulk (Tabula Muris, match FACS fractions) | 120 | B cells, T cells CD8, T cells CD4 conv |
| Petitprez-Sim | peritoneum, spleen, blood, tumor | mouse | pseudobulk (Tabula Muris, match FACS fractions) | 140 | B cells, NK cells, Tregs, T cells CD8, Macrophages |

| | | | | | |
|---|---|---|---|---|---|
| spillover-Sim | blood | human | pseudobulk (Hao, pure) | 550 | B cells, mDC, pDC, T cells CD4, T cells CD8, Treg, Monocytes, NK cells, ILC, platelets, Plasma cells |
| mRNAbias-Sim-blood | blood | human | pseudobulk (7 blood single-cell datasets, random) | 700 | B cells, mDC, pDC, T cells CD4, T cells CD8, Treg, Monocytes, NK cells, ILC, platelets, Plasma cells, Neutrophils, Erythrocytes |
| mRNAbias-Sim-Allen | brain | human | pseudobulk (Allen, random) | 100 | Astro, Endo, Excit, Inhib, Micro, Oligo, OPC |
| unknown-Sim | lung cancer | human | pseudobulk (Lambrechts, weighted) | 450 | B cells, T cells CD4, stromal cells, Macrophages, Tumor cells |
| resolution-Sim-Lambrechts | lung cancer and breast cancer | human | pseudobulk (Lambrechts, mirrorDB) | 50 | B cells, Macrophages alveolar, Macrophages, Monocytes classical, Monocytes non-classical, cDC1, cDC2, pDCs, T cells CD8 activated, T cells CD8 naive, T cells, D8 effector memory, T cells NK-like, T cells CD8 terminally exhausted, T cells CD4 non-regs, Tregs, NK cells |
| resolution-Sim-Wu | lung cancer and breast cancer | human | pseudobulk (Wu, mirrorDB) | 50 | B cells memory, B cells naive, CAFs MSC iCAF-like, CAFs myCAF-like, Macrophages, Monocytes, DCs, T cells CD8, T cells CD4, NK cells, NK T cells |
| resolution-Sim-Allen | brain | human | pseudobulk (Allen, mirrorDB) | 50 | Astro, Endo, L2/3 IT, L5 ET, L5 IT, L5/6 NP, L6 CT, L6 IT, L6 IT Car3, L6b, Lamp5, Micro-PVM, Oligo, OPC, Pvalb, Sncg, Sst Chodl, Vip, VLMC |
| ref-Sim-Lambrechts | lung cancer | human | pseudobulk (Lambrechts, mirrorDB) | 50 | B cells, NK cells, CD4 T cells, CD8 T cells, Tregs, Monocytes, Plasma cells |
| ref-Sim-Ma | lung | human | pseudobulk | 50 | B cells, NK cells, CD4 T cells, CD8 T cells, Tregs, |

| | | | | | |
|---|---|---|---|---|---|
| ynard | cancer | | (Maynard, mirrorDB) | | Monocytes, Plasma cells |

**Table S2**: list of all single-cell datasets used in this study.

| name | tissue | condition | organism | n_cells | n_samples | assay | cell types | reference |
|---|---|---|---|---|---|---|---|---|
| Hao | blood | healthy | human | 147391 | 8 | 10x 3' v3 | T cells CD4 conv, T cells CD8, ILC, Monocytes, NK cells, Plasma cells, Platelets, Tregs, mDC, B cells, pDC | https://doi.org/10.1016/j.cell.2021.04.048 |
| BioResource | blood | healthy | human | 101020 | 29 | 10x 3' | T cells CD4 conv, T cells CD8, ILC, Monocytes, NK cells, Plasma cells, Platelets, Tregs, mDC, B cells, pDC | https://doi.org/10.1038/s41591-021-01329-2 |
| Wu | breast | cancer | human | 88571 | 26 | 10x 3' and 5' v2 | B cells memory, B cells naive, CAFs MSC iCAF-like, CAFs myCAF-like, Macrophages, Monocytes, DCs, T cells CD8, T cells CD4, NK cells, NK T cells | https://doi.org/10.1038/s41588-021-00911-1 |
| Allen | brain | post-mortem | human | 76553 | 2 | SNARE-Seq 2 | Astro, Endo, L2/3 IT, L5 ET, L5 IT, L5/6 NP, L6 CT, L6 IT, L6 IT Car3, L6b, Lamp5, Micro-PVM, Oligo, OPC, Pvalb, Sncg, Sst, Sst Chodl, Vip, VLMC | https://doi.org/10.1038/s41586-021-03465-8 |
| TabulaSapiens | blood | healthy | human | 71057 | 9 | 10x 3' v3 | T cells CD4 conv, T cells CD8, ILC, Monocytes, NK cells, Plasma cells, Platelets, Tregs, mDC, B cells, pDC, Erythrocytes, Neutrophils | https://doi.org/10.1126/science.abl4896 |
| Lambrechts | lung | cancer | human | 64135 | 8 | 10x 3' v1 and v2 | Macrophages, T cells CD4 conv, Stromal cells, Tregs, Monocytes, NK cells, Tumor cells, mDCs, Mast cells, T cells CD8, Plasma cells, B cells, Epithelial cells, pDCs, Endothelial cells, Neutrophils | https://doi.org/10.1038/s41591-018-0096-5 |
| SchulteSchrepping | blood | healthy | human | 41378 | 21 | 10x 3' v2 | T cells CD4 conv, T cells CD8, ILC, Monocytes, NK cells, Plasma cells, Platelets, Tregs, mDC, B cells, pDC | https://doi.org/10.1016/j.isci.2021.103115 |
| Arunacha | blood | healthy | human | 24019 | 5 | 10x 3' | T cells CD4 conv, T cells CD8, ILC, | https://doi.org/10.10 |

| | | | | | | | | |
|---|---|---|---|---|---|---|---|---|
| lam | | | | | | v3 | Monocytes, NK cells, Plasma cells, Platelets, Tregs, mDC, B cells, pDC | 16/j.isci.2021.103115 |
| Heimlich | blood | healthy | human | 18930 | 7 | 10x 3' v3 | T cells CD4 conv, T cells CD8, Monocytes, NK cells, Platelets, Tregs, mDC, B cells, pDC | https://doi.org/10.1182/bloodadvances.2023011445 |
| DominguezConde | blood | healthy | human | 18766 | 2 | 10x 3' v3 | T cells CD4 conv, T cells CD8, ILC, Monocytes, NK cells, Plasma cells, Platelets, Tregs, mDC, B cells, pDC | https://doi.org/10.1126/science.abl5197 |
| Maynard | lung | cancer | human | 17546 | 29 | Smart-Seq2 | Monocytes, Macrophages, Endothelial cells, Mast cells, B cells, T cells CD8, T cells CD4 conv, Epithelial cell, Tumor cells, mDC, NK cells, Neutrophils, Tregs, Plasma cells, Stromal cells, pDC | https://doi.org/10.1016/j.cell.2020.07.017 |
| Lee | blood | healthy | human | 15512 | 4 | 10x 3' v3 | T cells CD4 conv, T cells CD8, ILC, Monocytes, NK cells, Plasma cells, Platelets, Tregs, mDC, B cells, pDC, Erythrocytes | https://doi.org/10.1126/sciimmunol.abd1554 |
| TabulaMuris | spleen | healthy | mouse | 9083 | 2 | Smart-Seq2 | B cells, NK cells, Tregs, T cells CD8, Macrophages, T cells CD4 conv, mDC | https://doi.org/10.1038/s41586-018-0590-4 |

**Table S3:** Cell type annotations for the Lambrechts, Wu and Allen datasets across the different resolution levels. For the Lambrechts dataset, the “fine” and “medium” levels were obtained with the dataset, while the “coarse” level was manually included. For the Wu dataset both the “fine” and “coarse” levels were included with the dataset, (‘celltype_major’ and ‘celltype_minor’). For the Allen dataset, only the fine level was included and the “coarse” level was manually prepared.

| **Fine annotation** | **Normal annotation** | **Coarse annotation** | **Dataset** |
|---|---|---|---|
| B cells | B cells | B cells | *Lambrechts* |
| Macrophages alveolar | Macrophages | Macrophages-Monocytes | *Lambrechts* |
| Macrophages | Macrophages | Macrophages-Monocytes | *Lambrechts* |
| Monocytes classical | Monocytes | Macrophages-Monocytes | *Lambrechts* |
| Monocytes non-classical | Monocytes | Macrophages-Monocytes | *Lambrechts* |
| cDC1 | mDCs | mDCs | *Lambrechts* |
| cDC2 | mDCs | mDCs | *Lambrechts* |
| pDCs | pDCs | pDCs | *Lambrechts* |
| T cells CD8 activated | T cells CD8 | T and NK cells | *Lambrechts* |
| T cells CD8 naive | T cells CD8 | T and NK cells | *Lambrechts* |
| T cells CD8 effector memory | T cells CD8 | T and NK cells | *Lambrechts* |
| T cells NK-like | T cells CD8 | T and NK cells | *Lambrechts* |
| T cells CD8 terminally exhausted | T cells CD8 | T and NK cells | *Lambrechts* |
| T cells CD4 non-regs | T cells CD4 | T and NK cells | *Lambrechts* |
| Tregs | T cells CD4 | T and NK cells | *Lambrechts* |
| NK cells | NK cells | T and NK cells | *Lambrechts* |
| B cells memory | - | B cells | *Wu* |
| B cells naive | - | B cells | *Wu* |
| CAFs MSC iCAF-like | - | CAFs | *Wu* |
| CAFs myCAF-like | - | CAFs | *Wu* |
| Macrophages | - | Myeloid | *Wu* |
| Monocytes | - | Myeloid | *Wu* |
| DCs | - | Myeloid | *Wu* |

| | | | |
|---|---|---|---|
| T cells CD8 | - | T cells | *Wu* |
| T cells CD4 | - | T cells | *Wu* |
| NK cells | - | T cells | *Wu* |
| NK T cells | - | T cells | *Wu* |
| Astro | - | Astro | *Allen* |
| Endo | - | Endo (endothelial cells) | *Allen* |
| L2/3 IT | - | Excit (excitatory neurons) | *Allen* |
| L5 ET | - | Excit (excitatory neurons) | *Allen* |
| L5 IT | - | Excit (excitatory neurons) | *Allen* |
| L5/6 NP | - | Excit (excitatory neurons) | *Allen* |
| L6 CT | - | Excit (excitatory neurons) | *Allen* |
| L6 IT | - | Excit (excitatory neurons) | *Allen* |
| L6 IT Car3 | - | Excit (excitatory neurons) | *Allen* |
| L6b | - | Excit (excitatory neurons) | *Allen* |
| Lamp5 | - | Inhib (Inhibitory neurons) | *Allen* |
| Micro-PVM | - | Micro (Microglia) | *Allen* |
| Oligo | - | Oligo (Oligodendrocytes) | *Allen* |
| OPC | - | OPC (Oligodendrocyte progenitor cells) | *Allen* |
| Pvalb | - | Inhib (Inhibitory neurons) | *Allen* |
| Sncg | - | Inhib (Inhibitory neurons) | *Allen* |
| Sst | | Inhib (Inhibitory neurons) | *Allen* |
| Sst Chodl | - | Inhib (Inhibitory neurons) | *Allen* |
| Vip | - | Inhib (Inhibitory neurons) | *Allen* |
| VLMC | - | Endo (endothelial cells) | *Allen* |
